# Kinetochore-mediated microtubule assembly and Augmin-dependent amplification drive k-fiber maturation in mammals

**DOI:** 10.1101/2021.08.18.456780

**Authors:** Ana C. Almeida, Joana Oliveira, Danica Drpic, Liam P. Cheeseman, Joana Damas, Harris A. Lewin, Denis M. Larkin, Paulo Aguiar, António J. Pereira, Helder Maiato

## Abstract

Chromosome segregation in mammals relies on the maturation of a thick bundle of kinetochore-attached microtubules known as k-fibers. How k-fibers mature from initial kinetochore-microtubule attachments remains a fundamental question. By combining functional perturbations in Indian muntjac, a placental mammal with the lowest known chromosome number (n=3) and distinctively large kinetochores, with fixed- and subsecond live-cell super-resolution CH-STED nanoscopy, we uncovered the mechanism by which Augmin mediates k-fiber maturation. Augmin promoted kinetochore microtubule turnover by sustaining microtubule formation from kinetochores and poleward flux, regardless of pioneer centrosomal microtubules. Tracking of microtubule growth events within individual k-fibers revealed a wide angular dispersion, consistent with Augmin-mediated branched microtubule nucleation. Augmin depletion reduced the frequency of microtubule growth events within k-fibers and hampered normal repair after acute k-fiber injury by laser microsurgery. Our work underscores the contribution of microtubule formation from kinetochores and Augmin-mediated microtubule amplification for k-fiber maturation and spindle assembly in mammals.

## Introduction

Accurate chromosome segregation during mitosis relies on the formation of a thick bundle of microtubules that attach at the kinetochore region of each chromosome to form kinetochore fibers (k-fibers) [1]. While the molecular basis of end-on kinetochore-microtubule attachments has been elucidated in recent years and was shown to involve the conserved Ndc80 complex [2], the mechanism by which mammalian kinetochores attach up to dozens of microtubules within a matter of minutes remains poorly understood. For years, this process, known as k-fiber maturation, was thought to rely on consecutive rounds of “search-and-capture” by centrosomal microtubules [3]. However, this proved to be highly inefficient [4] and inconsistent with the rapid acceleration of k-fiber maturation after a relatively slow initial microtubule capture rate at kinetochores [5]. Moreover, “search-and-capture” by centrosomal microtubules cannot explain k-fiber formation and maturation in cells that are naturally devoid of centrosomes, such as in land plants or female oocytes [6], or after experimental centrosome inactivation in animal somatic cells [7, 8].

Short non-centrosomal microtubules can be nucleated in the vicinity of chromosomes and kinetochores due to the microtubule stabilizing activity promoted by a Ran-GTP gradient and/or the chromosomal passenger complex [9–14]. These short microtubules are then captured and oriented with their plus-ends towards the kinetochore by CENP-E/kinesin-7 motors, and pre-formed k-fibers subsequently incorporated into the spindle by a Dynein-mediated interaction with non-kinetochore microtubules [14–19]. Augmin, an octameric Y-shaped complex that recruits γ-tubulin to pre-existing microtubules, triggers branched microtubule nucleation, thereby contributing to rapid microtubule amplification in the spindle [20–29]. In particular, the Augmin complex has been previously implicated in k-fiber formation [20, 22, 30, 31], but the underlying mechanism remains unclear. On one hand, Augmin was found to interact with Ndc80 and was shown to be required for kinetochore-driven microtubule formation in *Drosophila,* as well as centrosome-dependent microtubule nucleation in human cells [32, 33]. On the other hand, Augmin-mediated microtubule amplification was recently shown to be the predominant source of spindle microtubules in human somatic cells, and was proposed to account for the directional bias of microtubule growth towards the kinetochores after initial capture of pioneer centrosomal microtubules [34]. However, due to intrinsic limitations imposed by the high chromosome number and the sub-diffraction size of human kinetochores and associated k-fibers, Augmin’s role in k-fiber maturation has not been directly assessed. Moreover, the recent finding that most kinetochores in human cells develop their own k-fibers by “sorting” short randomly oriented non-centrosomal microtubules that appear in the immediate vicinity of the kinetochores [16], calls into question the requirement of pioneer centrosomal microtubules for k-fiber maturation.

The Indian muntjac (*Muntiacus muntjak)*, commonly known as “barking deer”, is a placental mammal whose females have the lowest known chromosome number of their class (n=3) [35]. As a result of repeated cycles of tandem and centric fusions [36, 37], Indian muntjac cells have long and morphologically distinct chromosomes with unusually large kinetochores (up to 2 μm linear length) that bind up to 60 microtubules [38–40]. These unique cytological features, combined with recent large-scale ruminant genome sequencing efforts [41], create the ideal conditions to directly dissect the molecular mechanism underlying k-fiber maturation in mammals. Here we used RNAi and high-resolution live-cell microscopy to investigate the role of 64 conserved mitotic proteins in mitotic spindle assembly and chromosome segregation in Indian muntjac cells. Assisted by unprecedented subsecond live-cell super-resolution CH-STED nanoscopy analysis [42] of microtubule growth within individual k-fibers and direct perturbation of k-fiber structure by laser microsurgery, we identified Augmin as the main driver of k-fiber maturation. This function was distinct from the role of the Ndc80 complex in the stabilization of end-on kinetochore-microtubule attachments and independent of pioneer centrosomal microtubule scaffolds. Instead, Augmin promoted microtubule growth from kinetochores and poleward flux. Most striking, a wide angular dispersion of microtubule growth events within individual k-fibers was observed, consistent with Augmin-mediated microtubule branching, but overlooked in current models of k-fiber formation and maturation. Altogether, our work directly elucidates how Augmin mediates k-fiber maturation in mammals and establishes the Indian muntjac as a powerful system for functional studies of mitosis.

## Results

### A live-cell RNAi screen in Indian muntjac fibroblasts identifies Augmin as a critical spindle assembly factor required for chromosome segregation

We used high-resolution live-cell microscopy combined with RNAi in hTERT-immortalized Indian muntjac fibroblasts [43] stably expressing histone H2B-GFP (to visualize chromosomes) and labelled spindle microtubules with 50 nM of SiR-tubulin [38, 44] to screen the roles of 64 conserved mitotic genes in spindle assembly and chromosome segregation (Fig. 1A, B). Control cells took 25 ± 8 minutes (mean ± standard deviation (S.D.)) from nuclear envelope breakdown (NEBD) until the completion of chromosome alignment (metaphase), or 37 ± 7 (mean ± S.D.) minutes until anaphase onset (AO) (Fig. 2A, A’, B and Fig. S1). Upon RNAi, phenotypical fingerprints were generated for each protein based on the fraction of cells that exhibited one or more of the following defects: A) incomplete congression and faster mitosis (NEBD-AO<23 minutes); B) incomplete congression and normal mitotic duration (23≤NEBD-AO<52 minutes); C) incomplete congression and prolonged mitosis (NEBD-AO≥52 minutes); D) congression delay (NEBD-metaphase≥41 minutes); E) metaphase delay (metaphase-AO≥28 minutes); F) anaphase lagging chromosomes; G) mitotic death and H) cytokinesis failure (Fig. 1A, Fig. 2A’ and Fig. S1). To facilitate the visualization of the observed phenotypes, we set up a public repository where time-lapse movies, phenotypical fingerprints and siRNA sequences for each gene can be conveniently browsed, and is freely available as a community resource (http://indianmuntjac.i3s.up.pt). An unbiased systematic cluster analysis defined ten distinct clusters and few “orphan” proteins that highlight hierarchical relationships based on phenotypic similarities and respective frequencies (Fig. 1B). Among others, depletion of the Ndc80 complex (Nuf2, Ndc80 and Spc24), Aurora A, chTOG or the chromosomal passenger complex (INCENP, Survivin and Aurora B) was highly detrimental for spindle assembly and/or chromosome segregation (Fig. 1B, Fig. 2A, A’, C and Fig. S1). Interestingly, co-depletion of VASH1 and VASH2, two recently identified carboxypeptidases involved in α-tubulin detyrosination [45, 46], clustered together with CENP-E/kinesin-7, providing genetic evidence for the recently proposed role of microtubule detyrosination in the regulation of CENP-E-dependent congression of pole-proximal chromosomes [47]. Surprisingly, depletion of HURP and TPX2, two proteins previously implicated in Ran-GTP-dependent acentrosomal k-fiber formation [13, 48–51] resulted only in very mild, if any, mitotic defects (Fig. 1B, Fig. 2A, A’, C, Fig. S1). In contrast, depletion of Eg5/kinesin-5 or the Augmin complex subunit HAUS6 emerged as the most deleterious conditions for mitosis in Indian muntjac fibroblasts (Fig. 1B, Fig. 2A, A’, C, Fig. S1). Because the critical role of Eg5/kinesin-5 motor activity in centrosome separation and bipolar spindle assembly is well established [52], we focused on dissecting the mechanism by which Augmin impacts spindle assembly and chromosome segregation.

**Fig. 1.**
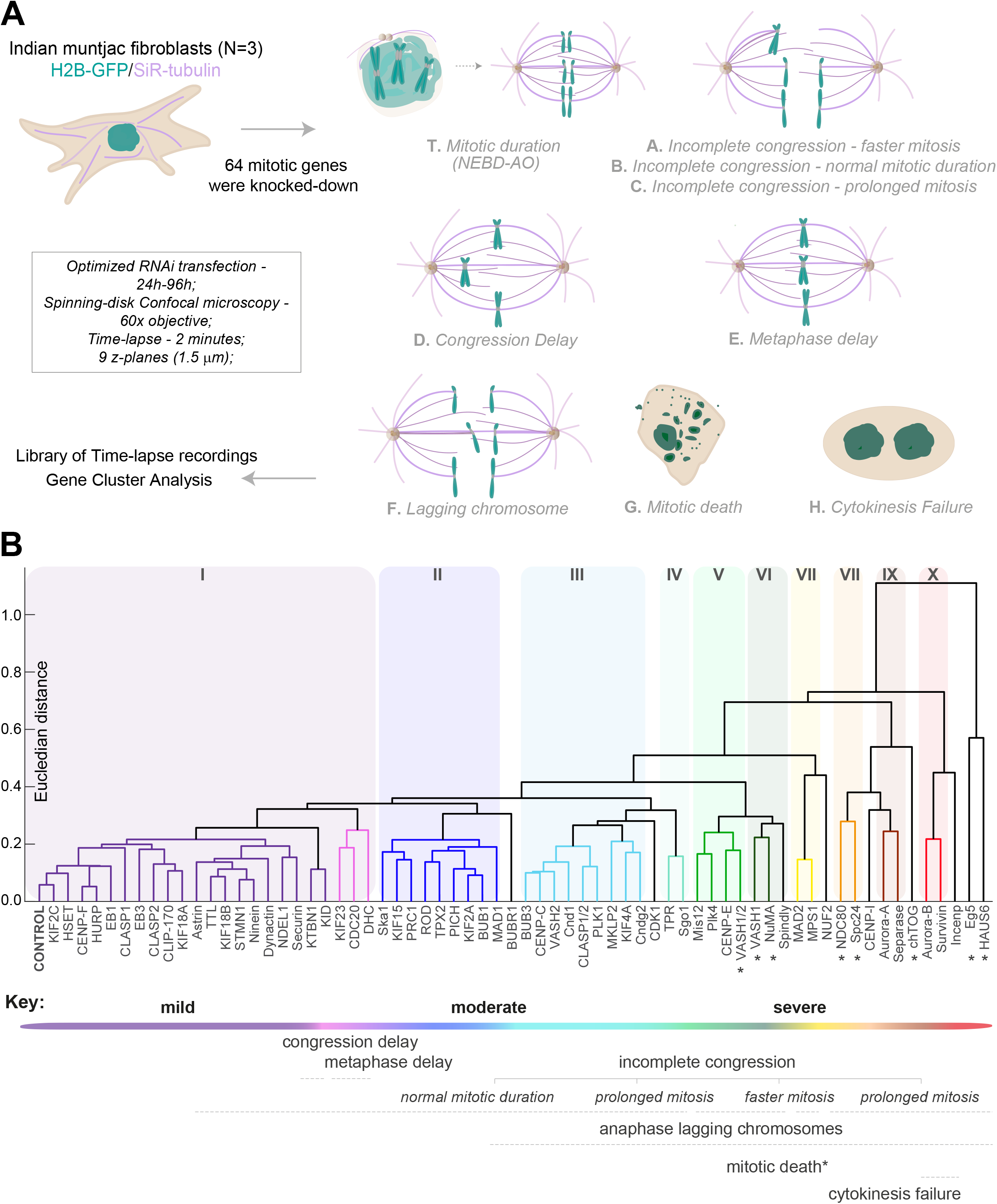
A live-cell RNAi screen in Indian muntjac fibroblasts identifies Augmin as a critical spindle assembly factor required for chromosome segregation. **A**, schematic representation of the mitotic screening performed in Indian muntjac fibroblasts stably expressing H2B-GFP to visualize chromosomes and labeled with 50 nM SiR-tubulin to image spindle microtubules. 64 mitotic genes were knocked down by RNAi for 24h, 48h, 72h or 96h, depending on the protein of interest. Mitotic timings (T. NEBD-AO) were determined and genes blindly clustered based on the probability of occurrence of eight binary features: A) incomplete congression and faster mitosis (NEBD-AO<23 minutes); B) incomplete chromosome and normal mitotic duration (23≤NEBD-AO<52 minutes); C) incomplete congression and prolonged mitosis (NEBD-AO≥52 minutes); D) congression delay (NEBD-metaphase≥41 minutes); E) metaphase delay (metaphase-AO≥28 minutes); F) anaphase lagging chromosomes; G) mitotic death and H) cytokinesis failure. **B,** dendrogram highlighting the hierarchical relationships between ten distinct clusters (I-X) and few “orphan” proteins based on phenotypic similarities and respective frequencies. The severity of the defects increased from left to right. Euclidean distance was used as the distance metric to compare the phenotypical fingerprints.

**Fig. 2.**
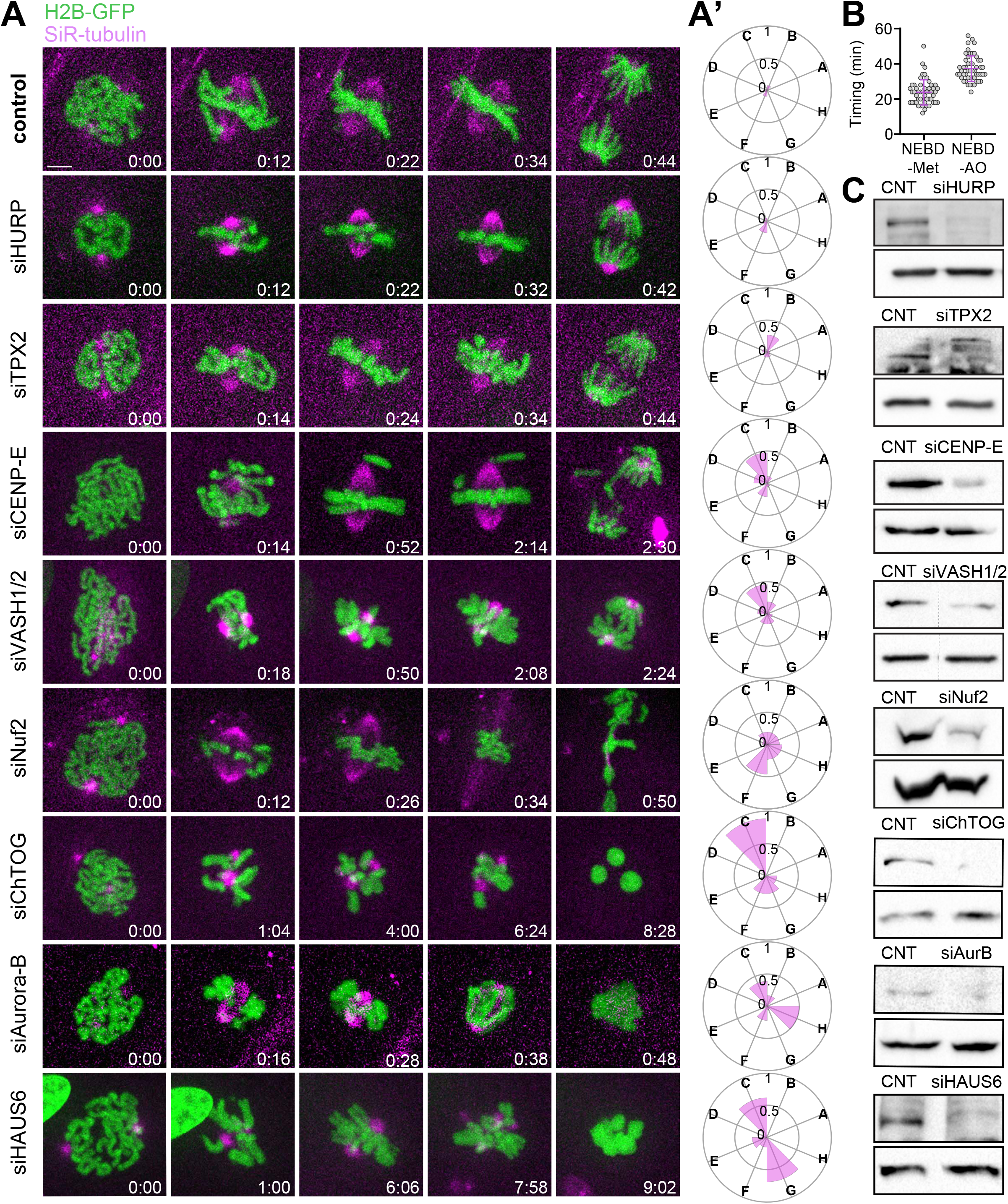
Augmin depletion emerged as one of the most deleterious conditions for mitosis in Indian muntjac fibroblasts. **A**, representative examples of the different protein clusters obtained by live-cell spinning-disk confocal microscopy of Indian muntjac fibroblasts stably expressing H2B-GFP (green) and treated with SiR-tubulin (magenta) after RNAi-mediated depletion: siHURP (n= 17 cells), siTPX2 (n= 26 cells), siCENP-E (n= 25 cells), siVASH1/2 (n= 26 cells), siNuf2 (n= 26 cells), siChTOG (n= 18 cells), siAurora B (n=18 cells) and siHAUS6 (n= 13 cells). Mock transfection (lipofectamine only) was used as control (n= 52 cells). Scale bar: 5 µm. Time: h:min. **A’,** radar plots illustrating the phenotypic fingerprints - probability of occurrence of the 8 analyzed features (A-G, see Fig. 1A) - for each silenced gene. Zero corresponds to a null-event and 1 to all cells displaying a certain event. **B,** control mitotic timings from NEBD-Metaphase (25 ± 8; mean ± S.D.) and NEBD-AO (37 ± 7; mean ± S.D.). **C,** protein lysates obtained after RNAi were immunoblotted with an antibody specific for each protein of interest (upper band), except for VASH1/2, where only anti-VASH1 was used and Nuf2, where anti-Hec1 was used. The bottom band corresponds to anti-GAPDH (siTPX2, siMad2, siVASH1/2, siNuf2 and siHAUS6), anti-α-tubulin (siHURP and siAuroraB) or anti-vinculin (siCENP-E and siChTOG), which were used as loading controls. CNT = control.

### Augmin recruits γ-tubulin to the spindle region and promotes robust k-fiber formation in Indian muntjac fibroblasts

We started by using conventional fluorescence microscopy in fixed cells to validate whether Augmin’s requirement for k-fiber formation was conserved in Indian muntjac fibroblasts. To standardize conditions and allow enough time for spindle assembly, both control and HAUS6-depleted cells were arrested in mitosis for 1.5 hours with the proteasome inhibitor MG-132. In agreement with our live-cell data, mitotic spindle length after HAUS6 depletion was reduced almost 50% relative to controls (Fig. 2A, A’, Fig. S2A, A’). In line with previous findings in *Drosophila* and human cells [20, 21, 29, 31], HAUS6 was found associated with spindle microtubules in Indian muntjac fibroblasts (Fig. S3A), and its depletion drastically reduced γ-tubulin accumulation in the spindle region (Fig. S2B, B’). These phenotypes were the specific result of Augmin perturbation since RNAi-mediated depletion of another Augmin subunit (HAUS1) was indistinguishable from HAUS6 depletion (Fig. S3B). Immunofluorescence analysis with antibodies against Mad2, which accumulates at unattached kinetochores [53], and HURP, which decorates the kinetochore-proximal ends of k-fibers [48], revealed that robust k-fiber formation was significantly compromised in HAUS6-depleted cells (Fig. S2C, C’, D, D’). Indeed, cold treatment at 4°C for 5 minutes to selectively destabilize non-kinetochore microtubules in the spindle confirmed that k-fibers were nearly absent after HAUS6 depletion (Fig. S2E, E’). Depletion of Ndc80, an outer kinetochore protein required for the stabilization of end-on microtubule attachments [2] was used as a positive control (Fig. S2D, D’, E, E’, Fig. S3C, D).

To gain additional insight into the role of Augmin in k-fiber formation, we optimized fixation conditions to preserve microtubule structure (see Materials and Methods) and inspected HAUS6-depleted cells by super-resolution coherent-hybrid stimulated emission depletion (CH-STED) nanoscopy, which markedly improves contrast in complex 3D objects like the mitotic spindle relative to conventional 2D-STED [42]. This analysis confirmed the absence of robust k-fibers after Augmin perturbation (Fig. 3A). Surprisingly, we found that HAUS6-depleted cells exhibited overly elongated astral microtubules (Fig. 3A, B). To rule out a possible role for the Augmin complex in centrosome-dependent microtubule nucleation [33], we performed a microtubule regrowth assay after treatment with the microtubule-depolymerizing drug nocodazole for 2 hours, a condition that completely depolymerized all microtubules (Fig. S4A and Fig. S5A), followed by nocodazole washout and fixation after 2, 5 and 10 minutes, in the presence or absence of HAUS6 (Fig. S4A, B). We found that after HAUS6 depletion, centrosome-nucleated astral microtubules grew significantly longer than controls, 5 and 10 minutes after nocodazole washout, despite being slightly shorter at 2 minutes (Fig. S4A, B). By comparison, depletion of Ndc80, led to a similar, yet less pronounced, outcome (Fig. S4A, B). In contrast, perturbation of the TOG-domain proteins chTOG or CLASPs, which promote microtubule polymerization [54], significantly compromised microtubule regrowth from centrosomes after nocodazole treatment/washout in all time points (Fig. S4A, B). These results strongly suggest that, regardless of the underlying molecular nature, experimental perturbation of k-fiber formation in Indian muntjac cells is sufficient to bias tubulin polymerization towards astral microtubules.

**Fig. 3.**
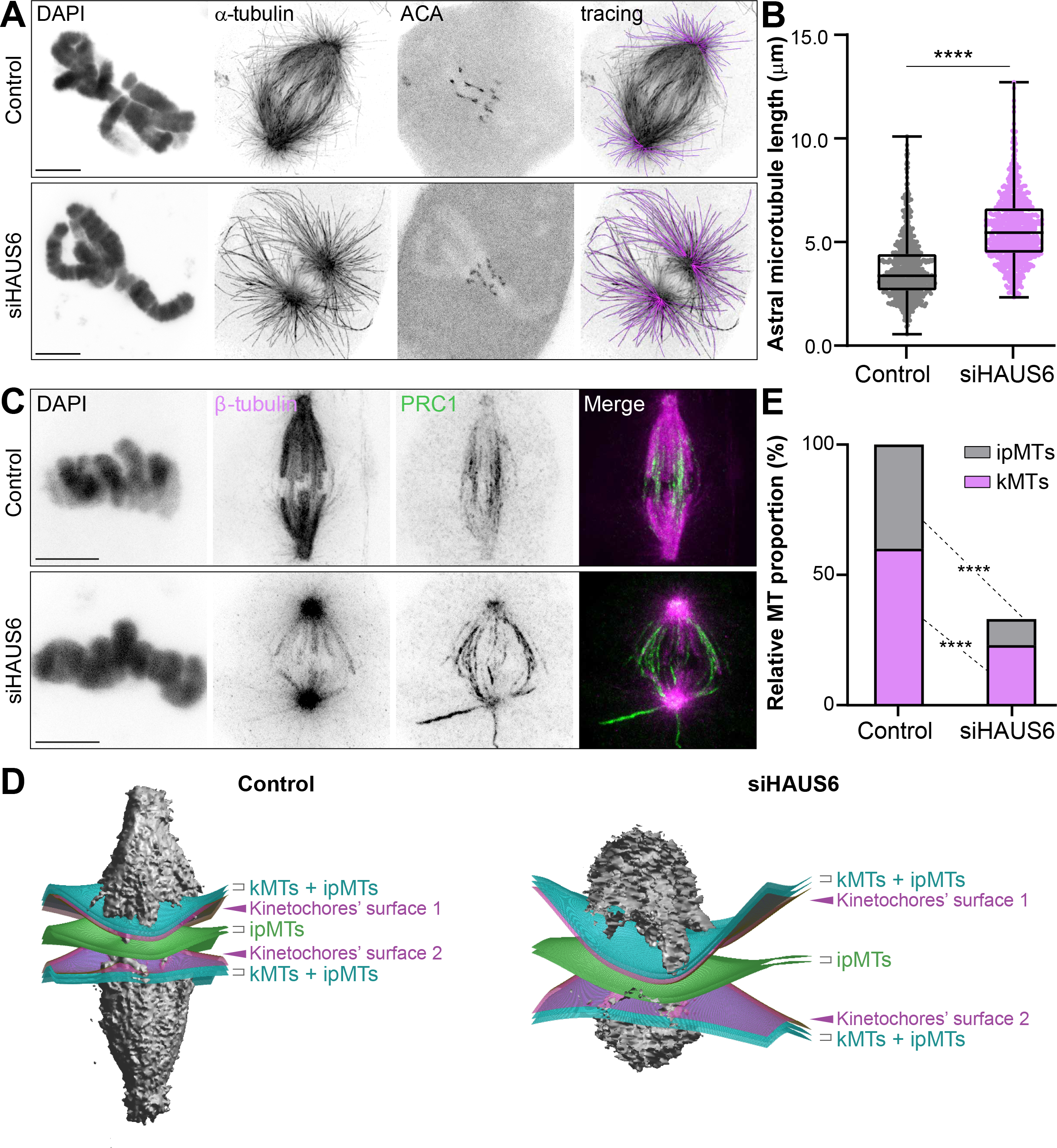
Augmin contributes to k-fiber and interpolar microtubule formation. Representative images of control and HAUS6-depleted cells acquired by CH-STED nanoscopy. **A,** DAPI, α-tubulin and ACA are shown in inverted grayscale. Astral microtubule tracks are represented in magenta (tracing). **B**, quantification of astral microtubule length (n= 409 Control astral microtubules; n= 324 siHAUS6 astral microtubules). The box plot determines the interquartile range; the line inside the box represents the median. **C,** immunofluorescence of Indian muntjac fibroblasts with β-tubulin (magenta) and PRC1 (green) antibodies. **D,** representative 3D representations of mitotic spindles in control and HAUS6-depleted cells, illustrating kinetochore surfaces’ 1 and 2 (magenta), as well as the plates that define the measurement volumes, corresponding to interpolar microtubules (ipMTs, green) and k-fibers (kMTs) plus ipMTs (cyan). **E,** quantification of ipMTs and kMTs in control and HAUS6-depleted cells. Proportion relative to control levels is represented for HAUS6-depleted cells (n= 14 Control cells; n= 12 siHAUS6 cells). ****p≤0.0001. Scale bar: 5 µm.

To obtain a quantitative picture of Augmin’s contribution to k-fiber and interpolar microtubule formation, we processed Indian muntjac fibroblasts for immunofluorescence with antibodies against PRC1 and β-tubulin. PRC1 was enriched along overlapping interpolar microtubules (also known as bridging fibers) [55] in control metaphase cells (Fig. 3C). Surprisingly, HAUS6 depletion caused the dispersion of PRC1 to both parallel and anti-parallel microtubules (Fig. 3C). For this reason, we implemented a quantitative assay to determine the proportion of kinetochore and non-kinetochore microtubules relying exclusively on the β-tubulin signal. Based on the coordinates of k-fiber endpoints, a 200 nm-thick curved plate (a spline surface) that smoothly crossed all inter-kinetochore midpoints (i.e. centromeres) along the metaphase plate was generated and assumed to be crossed only by interpolar microtubules. A second spline plate was defined adjacent to the kinetochores but shifted polewards by 100 nm. This plate was crossed by all interpolar microtubules and k-fibers in the corresponding half-spindle (Fig. 3D). This analysis revealed that HAUS6 depletion caused ∼60% reduction in the total spindle microtubule population, affecting both kinetochore and interpolar microtubules (Fig. 3E). Taken together, these data demonstrate that Augmin recruits γ-tubulin to the spindle and is required for k-fiber and interpolar microtubule formation in Indian muntjac fibroblasts.

### Augmin sustains centrosome-independent microtubule growth from kinetochores

The roles of centrosome- and kinetochore-dependent pathways in k-fiber formation have long been recognized [56], but their relative contributions remain unclear. For instance, centrosome-nucleated astral microtubules may be captured by chromosomes at their kinetochores [57, 58], but recent correlative light and electron microscopy studies in early prometaphase in human cells revealed that most k-fibers form by capturing short randomly oriented non-centrosomal microtubules that appear in the immediate vicinity of the kinetochores [16], suggesting that the kinetochore pathway is the predominant mechanism accounting for k-fiber formation in mammals. We therefore sought to investigate the role of Augmin in centrosome-independent, kinetochore-driven, k-fiber formation. To do so in a systematic way, we followed microtubule regrowth from Indian muntjac kinetochores after nocodazole treatment/washout, which recapitulates microtubule growth from kinetochores under physiological conditions [12, 13, 16], after treating cells with centrinone, a Plk4 inhibitor that prevents centriole duplication [59] (Fig. 4A, A’). Successful elimination of centrioles was confirmed by the loss of GFP-Centrin-1 by CH-STED microscopy (Fig. S5B). We found that short microtubule stubs appeared virtually in all kinetochores in control, HAUS6- and Ndc80-depleted cells (Fig. 4A-C, Fig. S5C, D), suggesting that the Augmin and Ndc80 complexes are dispensable for the initial step of microtubule nucleation in the vicinity of kinetochores in mammals. However, both HAUS6- and Ndc80-depleted cells showed a significant decrease in the fraction of kinetochores that remained associated with microtubules over time (Fig. 4A, B, Fig. S5C, D). Importantly, whereas HAUS6 depletion prevented kinetochore microtubules from growing to the same extent as in controls, Ndc80 depletion led to longer microtubules that appeared to associate laterally with kinetochores (Fig. 4A, C, Fig. S5C, E). These results suggest that while Ndc80 is necessary to stabilize end-on microtubule attachments after nucleation in the vicinity of kinetochores, Augmin is required to sustain microtubule growth from kinetochores.

**Fig. 4.**
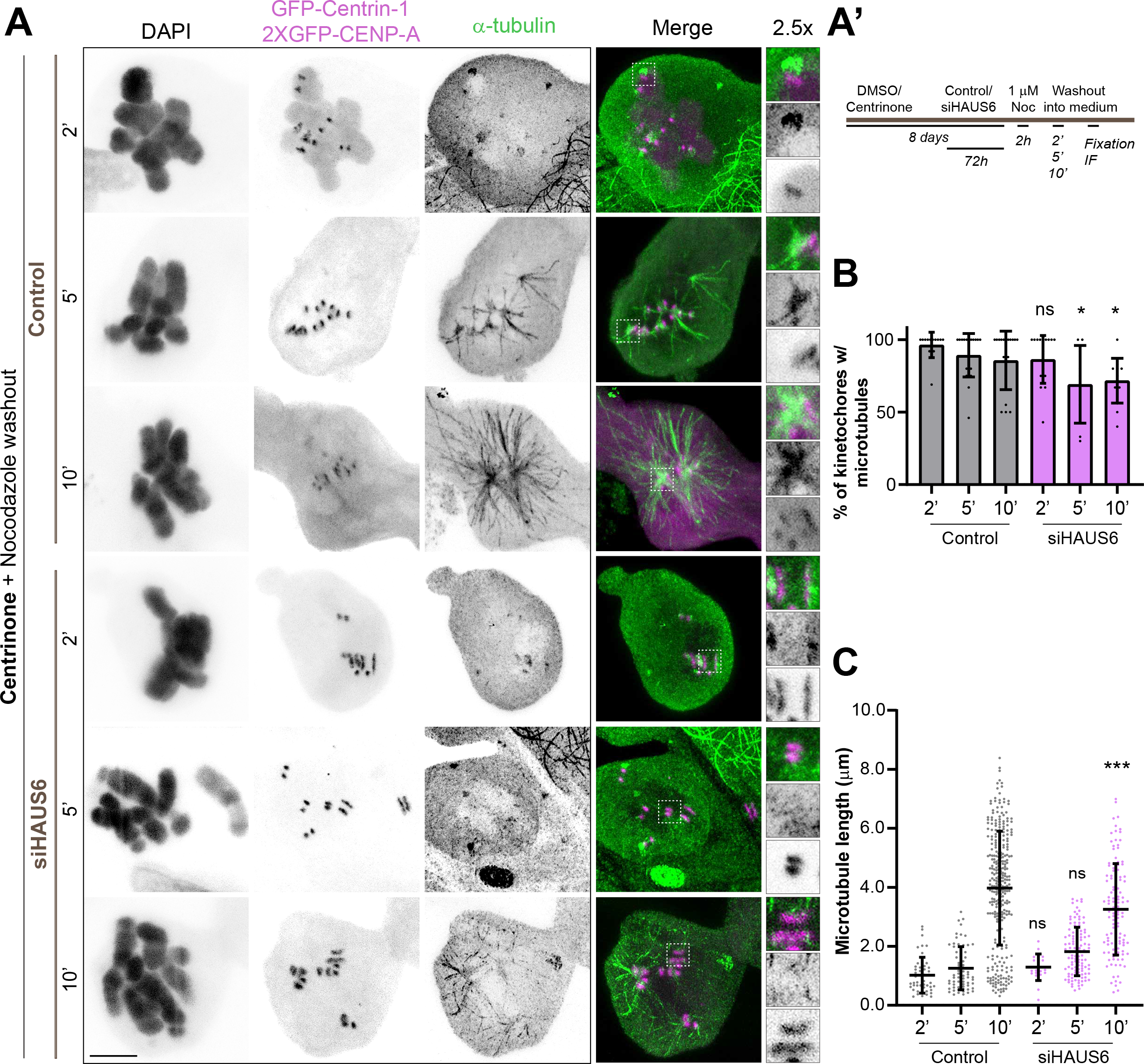
Augmin sustains centrosome-independent microtubule growth from kinetochores. **A**, representative CH-STED images of Indian muntjac cells stably expressing GFP-Centrin-1 (magenta) and 2xGFP-CENP-A (magenta) treated with centrinone/DMSO for 8 days and transfected with HAUS6 RNAi for 72 hours. A microtubule regrowth assay at 37 °C was performed. Cells were treated with the microtubule-depolymerizing drug nocodazole for 2 hours, followed by drug washout and fixation after 2, 5 and 10 minutes. Cells were immunostained with anti-α-tubulin antibody (green) and DAPI (inverted grey scale). Insets show 2.5x magnification of selected regions with kinetochore and nucleated microtubules (grayscale for single channels of CENP-A and α-tubulin). Experimental setup is described in **A’**. Percentage of kinetochores with microtubules and overall microtubule length are shown in **B** and **C**, respectively (Control 2’: n= 14 cells/49 microtubules, Control 5’: n= 20 cells/66 microtubules, Control 10’: n= 19 cells/313 microtubules; siHAUS6 2’: n= 20 cells/20 microtubules; siHAUS6 5’: n= 9 cells/111 microtubules; siHAUS6 10’: n= 14 cells/119 microtubules). Mean ± S.D.; ns: not significant, *p≤0.05, ***p≤0.001. Scale bars: 5 µm.

### Augmin promotes kinetochore microtubule turnover and poleward flux

To investigate how Augmin sustains microtubule growth from kinetochores we started by implementing a live-cell CH-STED nanoscopy assay in Indian muntjac fibroblasts stably expressing GFP-CENP-A to visualize kinetochores, and Halo-tagged EB3 conjugated with the bright, photostable, far-red ligand JF646 [60, 61] to track growing microtubule plus ends for 2 minutes at 8 seconds and 100 nm resolution (Fig. 5A, A’, Fig. S6A). Quantitative analyses revealed that the velocity of poleward and anti-poleward chromosome movement, chromosome oscillation amplitude, but not oscillation period, were severely reduced upon HAUS6 or Ndc80 depletion (Fig. 5B-E). Moreover, while in control cells EB3 accumulated at kinetochores for one half period of chromosome oscillations corresponding to anti-poleward movement, EB3 only rarely associated with kinetochores after HAUS6 or Ndc80 depletion (Fig. 5F). Importantly, HAUS6, but not Ndc80, depletion led to a shorter kinetochore-to-pole distance (Fig. 5G), suggesting a distinct mode of action by which these proteins contribute to k-fiber formation and dynamics.

**Fig. 5.**
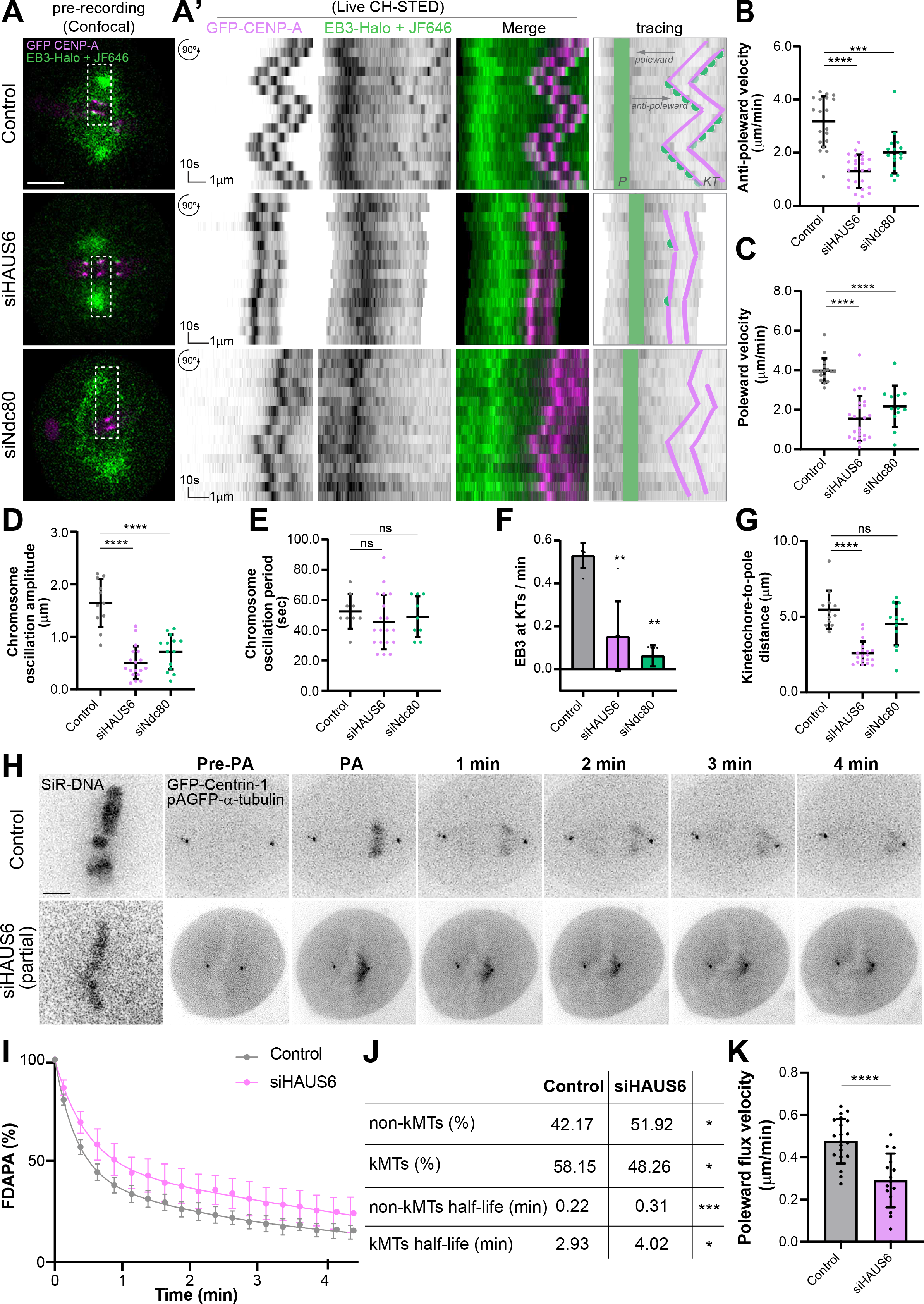
Augmin promotes kinetochore microtubule turnover and poleward flux. **A**, pre-recording snapshots of control, HAUS6- and Ndc80-depleted Indian muntjac fibroblasts stably expressing GFP-CENP-A (magenta) and EB3-Halo tag, conjugated with a far-red dye JF646 (green), imaged by confocal microscopy. **A’,** collapsed kymographs of live CH-STED recordings (time-lapse: 8 sec; pixel size: 40 nm). Graphical sketches on the right highlight chromosome movement over time (tracing); P-pole (green); KT-kinetochore (magenta); EB3 accumulation at kinetochore is shown in green. Quantitative analysis of chromosome anti-poleward, **B,** and poleward, **C**, velocities, chromosome oscillatory amplitude **D,** and period, **E**. Fraction of EB3 accumulation at kinetochore per minute (approximately one period) was measured from track data in **F,** and kinetochore-to-pole distance determined in **G**. Each data point represents one measurement. Horizontal bar: 1 µm; vertical bar: 10 s; (n=8 Control cells, n=9 siHAUS6 cells and n=8 siNdc80 cells). **H,** representative examples of control and partial HAUS6-depleted metaphase cells displaying photoactivatable pAGFP-α-tubulin (inverted grayscale), GFP-Centrin-1 (inverted grayscale) and labeled with 50 nM SiR-DNA to visualize chromosomes (inverted grayscale). “Pre-PA”: frame immediately before photoactivation; “PA”: frame immediately after photoactivation. **I,** normalized fluorescence dissipation after photoactivation (FDAPA) curves of control and partial HAUS6-depleted cells. Whole lines show double exponential curve fittings (R^2^>0.98) and error bars show 95% confidence interval for each time point. **J,** table showing the calculated microtubule percentages and turnover values for control and partial HAUS6-depleted cells (n= 20 Control cells; n= 11 siHAUS6 cells). **K,** microtubule flux velocity (n= 21 Control cells; n= 16 siHAUS6 cells). Each data point represents one measurement; mean ± S.D.; ns: not significant, *p≤0.05, **p≤0.01, ***p<0.001, ****p≤0.0001. Scale bar: 5 µm.

To determine whether Augmin is required to promote kinetochore microtubule turnover we used fluorescence dissipation after photoactivation (FDAPA) of GFP-α-tubulin [62] (Fig. 5H). By fitting the fluorescence decay over time to a double exponential curve (R^2^>0.98), we differentiated two spindle microtubule populations with fast and slow fluorescence decay (Fig. 5I) that have been attributed to less stable non-kinetochore microtubules and more stable kinetochore microtubules, respectively [62, 63]. We found that partial HAUS6 depletion by RNAi (note that optimal HAUS6 depletion completely disrupts k-fiber formation) significantly increased the half-life of both kinetochore and non-kinetochore microtubules (Fig. 5J). In parallel, by measuring the velocity by which the photoactivation mark on spindle microtubules moved relative to the metaphase plate (i.e. underwent poleward flux) [64], we found that it was reduced by ∼40% after partial HAUS6 depletion (Fig. 5K). Overall, these data indicate that Augmin promotes kinetochore and non-kinetochore microtubule turnover, while assisting poleward flux in metaphase cells.

### Microtubule growth within k-fibers show a wide angular dispersion and requires Augmin

Comparative fixed-cell super-resolution imaging revealed striking differences in k-fiber structure between control, HAUS6- and Ndc80-depleted cells, whereas depletion of TPX2, HURP, chTOG and CLASP1/2 did not compromise the formation of robust k-fibers, despite an obvious reduction in k-fiber length after chTOG or CLASP1/2 depletion (Fig. 6A, Fig. S6B). To directly investigate how Augmin mediates kinetochore microtubule turnover, we tracked microtubule growth events within a single k-fiber using live-cell CH-STED nanoscopy of EB3 comets in the kinetochore vicinity, now for 1 minute at 750 milliseconds and 100 nm resolution (Fig. 6B, Fig. S6A). These imaging conditions did not result in any obvious phototoxicity or relevant photobleaching throughout the recordings (Fig. S6C, D). Temporal projections of EB3 comets over consecutive frames in control cells revealed several microtubule growth events on a single k-fiber (Fig. 6B). Cross-correlation analysis of EB3 comets (see Materials and Methods) in control cells revealed a microtubule growth velocity of ∼9 μm/min, with an absolute angular dispersion of 37° ± 13° (mean ± S.D.) relative to the respective k-fiber axis perpendicular to the kinetochore plate (Fig. 6B, C, Fig. S6E). Remarkably, detailed inspection of collapsed kymographs within individual k-fibers allowed the direct visualization and discrimination of microtubule growth events that terminate or pass by the kinetochore (Fig. 6D). We found that HAUS6 or Ndc80 depletion caused a 30-40% reduction in the frequency of microtubule growth events that terminate at the kinetochore (Fig. 6D, E), with a corresponding reduction in inter-kinetochore distances (Fig. 6D, F), suggestive of compromised k-fiber formation. However, while HAUS6 depletion did not significantly affect the frequency of microtubule growth events that pass by the kinetochore, this was largely increased after Ndc80 depletion, consistent with a role of Ndc80 in the stabilization of end-on kinetochore-microtubule attachments. Of note, none of these experimental conditions altered kinetochore and non-kinetochore microtubule plus-end growth velocity, as determined by measuring the respective slopes from EB3 tracks on the kymographs (Fig. S6D), in line with our previous cross-correlation analysis (Fig. S6E). Overall, these data indicate that Augmin and Ndc80 mediate k-fiber formation by distinct mechanisms and directly demonstrates a role for Augmin in microtubule nucleation within k-fibers.

**Fig. 6.**
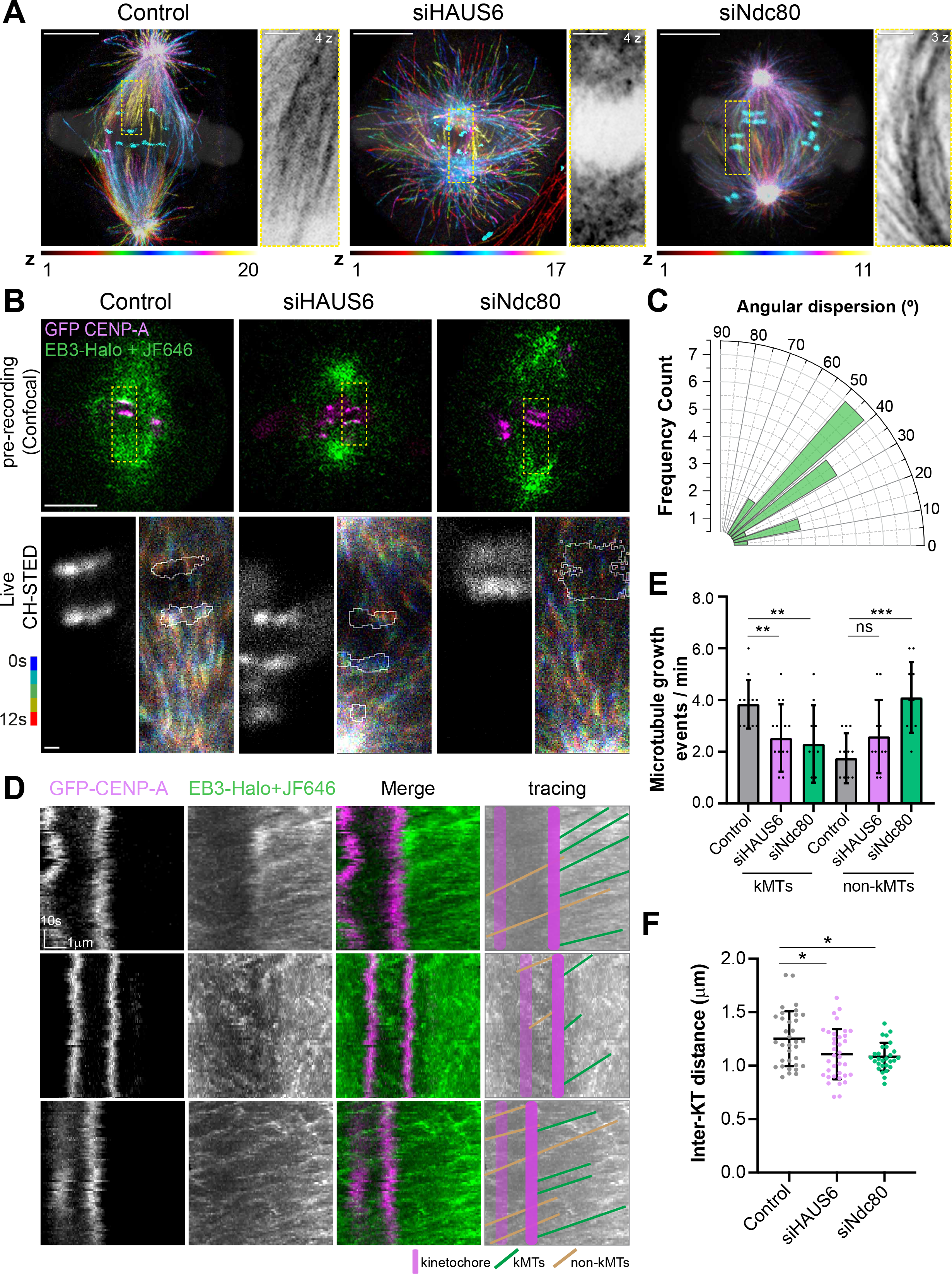
Microtubule growth within individual k-fibers show a wide angular dispersion and requires Augmin. **A**, representative super-resolution images of control, HAUS6- and Ndc80-depleted cells stained with α-tubulin, ACA (cyan) and DAPI (white; opacity 15%). Temporal color code tool on Fiji was used to match each α-tubulin z-plane to a different color. Scale bar: 5 µm. Insets show the maximum-intensity projection (selected z-planes) of a k-fiber (α-tubulin, inverted grayscale). Scale bar: 1 µm. **B,** Indian muntjac fibroblasts stably expressing GFP-CENP-A (magenta) and EB3-Halo tag, conjugated with a far-red dye JF646 (green) were used to track microtubule polymerization events within one k-fiber by live CH-STED microscopy (time-lapse: 750 ms; pixel size: 40 nm). Images on the left show a pre-recording snapshot (confocal). Images to the right show chromo-projections of the time-lapse movie of fluorescently labeled EB3 over time (right) and CENP-A tracing (white line and middle panel). A limited time-window of 12 sec (5 frames) was selected, allowing a fine time-discrimination of microtubule growing events. Scale bar: 500 nm. **C,** frequency count of EB3 comets’ angular dispersion relative to the k-fiber axis in control cells (n= 17 cells). **D**, corresponding collapsed kymographs of control, HAUS6- and Ndc80-depleted cells. Graphical sketches on the right highlight detected EB3 comets’ trajectories; KT-kinetochore (magenta); kMTs - green; non-kMTs – light brown. **E,** number of EB3 growing events per kinetochore (Control: n= 46 k-fibers comets/n= 21 interpolar comets/n= 12 cells; siHAUS6: n= 38 k-fibers comets/n= 41 interpolar comets/n= 15 cells; siNdc80 n= 23 k-fibers comets/n= 41 interpolar comets/n= 10 cells). **F,** inter-kinetochore distances were determined by measuring the distance between kinetochore pairs upon stable expression of GFP-CENP-A (Control n= 34 cells; siHAUS6 n= 36 cells; siNdc80 n= 28 cells). Mean ± S.D.; ns: not significant, *p≤0.05, **p≤0.01, ***p<0.001. Vertical bar: 10 sec; horizontal bar: 1 µm.

### Augmin is required for microtubule amplification from pre-existing kinetochore microtubules

To directly test whether Augmin is required for microtubule amplification from pre-existing kinetochore microtubules, we developed a laser microsurgery-based k-fiber maturation assay (Fig. 7A). In this assay, we used live Indian muntjac fibroblasts stably expressing GFP-α-tubulin and CENP-A fused with the bright, red-shifted, monomeric fluorescent protein mScarlet to visualize and discriminate mature k-fibers from other spindle microtubules by spinning-disk confocal microscopy (Fig. 7B). Then, we used a pulsed laser microbeam to acutely induce partial k-fiber damage and measured the respective kinetics of fluorescence recovery after surgery (FRAS) as a proxy for k-fiber recovery, in controls and after partial HAUS6 depletion by RNAi (Fig. 7A-C). Partial k-fiber damage was confirmed by correlative live-cell spinning-disk confocal microscopy and super-resolution CH-STED nanoscopy after fixation of the same cell, and the lack of the typical snap observed immediately after complete k-fiber severing, which results in two independent kinetochore- and pole-proximal microtubule stubs (Fig. S7A-C). A 300 nm spacer between the kinetochore-proximal stub and the ablated region was defined to exclude the contribution of microtubule poleward flux for k-fiber recovery (see Materials and Methods). Strikingly, we found that partially-ablated k-fibers in control cells took on average 9 seconds to recover 50% of the fluorescence intensity in the damaged region, corresponding to half the time required in HAUS6-depleted cells (Fig. 7B, C). Moreover, while all control cells recovered completely from partial k-fiber ablation within the first 30 seconds after surgery, partially HAUS6-depleted cells recovered only 80% (Fig. 7D). Overall, these data directly demonstrate that Augmin is required for microtubule amplification from pre-existing kinetochore microtubules, consistent with a critical role in k-fiber maturation.

**Fig. 7.**
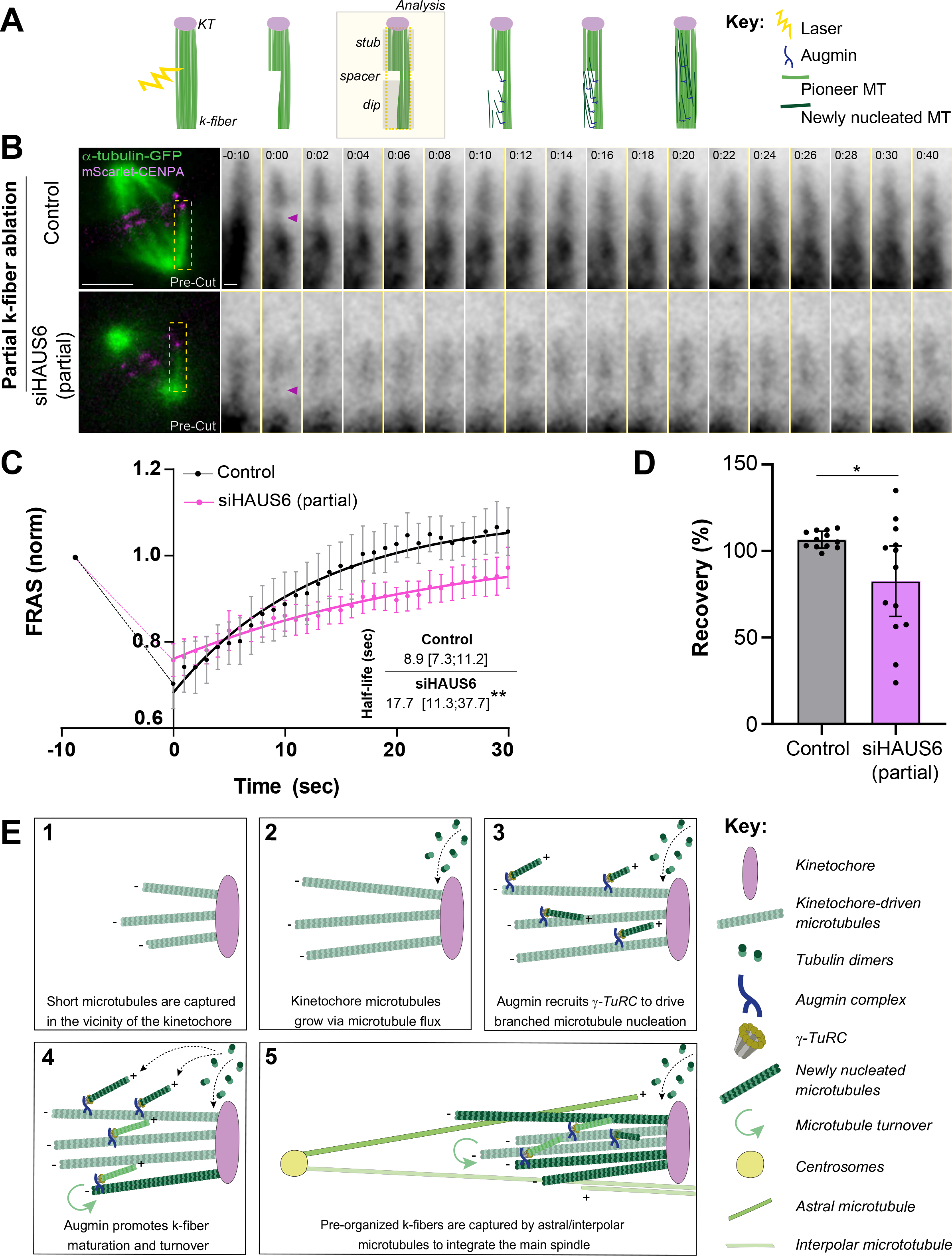
Augmin is required for microtubule amplification from pre-existing kinetochore microtubules. **A**, schematic summary of the laser microsurgery-based k-fiber injury/repair assay in control cells. Three ROI’s were defined for analysis (see Materials and Methods). **B,** representative spinning-disk confocal images of control and partial HAUS6-depleted Indian muntjac fibroblasts stably expressing GFP-α-tubulin (green) and mScarlet-CENP-A (magenta), illustrating microtubule recovery after partial k-fiber laser ablation. Schematic representation of the assay in control cells. Images on the left show the frame before the injury (Pre-Cut). Yellow dashed-rectangle indicates the ablated k-fiber. Scale bar: 5 µm. Insets show the analyzed k-fibers (GFP-α-tubulin, inverted grayscale). -10 sec represents arbitrary time before partial k-fiber ablation. Purple arrowhead points to the ablated k-fiber portion at time zero (first frame after laser ablation). Scale bar: 1 µm, time: min:sec. **C,** kinetics of fluorescence recovery after surgery (FRAS) was determined as a proxy for k-fiber recovery, in controls and after partial HAUS6-depletion by RNAi. Whole lines show single exponential fitting curve. Each data point represents the mean ± 95% confidence interval; **p≤0.01. **D,** percentage of fluorescence recovery from partial k-fiber ablation within the first 30 seconds after surgery relative to fluorescence decrease at time=0, in control and partial HAUS6-depleted cells (n= 12 Control cells; n= 13 siHAUS6 cells). Mean ± sd; *p≤0.05; **p≤0.01. **E,** proposed model for mammalian k-fiber maturation by Augmin: 1) initial microtubule organization and capture in the vicinity of kinetochores; 2) microtubule growth from the kinetochores via microtubule flux; 3) Augmin recruits γ-TuRCs to drive branched microtubule nucleation; 4) microtubule amplification by Augmin promotes k-fiber maturation and turnover; 5) pre-organized k-fibers are captured by astral and/or interpolar microtubules to integrate the main spindle.

## Discussion

Here we implemented the simplest mammalian model system known to date based on female Indian muntjac fibroblasts with just three pairs of chromosomes for the molecular dissection of mitosis. The rationale behind this approach was that a simpler mammalian system would reveal previously overlooked molecular requirements upon genetic perturbation and detailed phenotypic analysis. In particular, the unique cytological features of Indian muntjac cells offer the opportunity to directly address fundamental questions related with kinetochore biology and function that are not possible in any other system, including human cells. This is the case of k-fiber maturation, whose underlying molecular mechanism remains poorly understood. Based on its established roles in microtubule amplification from pre-existing microtubules, both *in vitro* and in cells, including in the mitotic spindle [25-27, 65, 66], the Augmin complex is a strong candidate to mediate k-fiber maturation in mammals [67, 68]. However, the underlying mechanism remained unclear due to two main reasons. First, Augmin subunits were found to interact with the Ndc80 complex [32, 33], which is required for the stabilization of end-on kinetochore-microtubule attachments, offering an alternative mechanistic explanation that is independent of Augmin’s roles in microtubule amplification from pre-existing microtubules. Second, the intrinsic limitations of human kinetochores and k-fibers, whose structures cannot be resolved by conventional light microscopy, precluded the experimental dissection of the process. By comparing the requirements of the Augmin and Ndc80 complexes, among other candidate proteins, for k-fiber formation in Indian muntjac cells, supported by subsecond live-cell super-resolution nanoscopy of microtubule growth events on individual k-fibers and acute laser microsurgery-based perturbations, our work directly investigates the mechanism by which Augmin promotes k-fiber maturation in mammals. We found that Augmin’s role in this process is clearly distinct from that of Ndc80 and is consistent with branched microtubule nucleation from pre-existing kinetochore microtubules. Indeed, Augmin was required to recruit γ-tubulin to the spindle and to sustain, but not to initiate, centrosome-independent microtubule growth from kinetochores. This contrasts with the previous implication of Augmin in the nucleation and/or initial stabilization of chromosome-induced microtubules in *Drosophila* cells [32], but is in agreement with previous works in several mammalian cells, including human, that showed that the vast majority of spindle microtubules do not have a centrosomal origin [7, 34, 69]. Importantly, with no prejudice to the role of pioneer centrosomal microtubules that might assist the subsequent microtubule amplification cascade by Augmin [34], our data suggest that Augmin contributes to k-fiber maturation even in the absence of pre-existing centrosomal microtubules. This is consistent with the fact that functional spindles are able to assemble in several animal species, including humans, after perturbation of centrosome function [7, 8, 69–74]. In these cases, short microtubules nucleated in the vicinity of and subsequently oriented and captured by kinetochores [14, 16] might work as amplification platforms for centrosome-independent formation of k-fibers, while still ensuring the directional bias of microtubule growth towards the kinetochore. When centrosomes are present, Augmin-dependent microtubule amplification of short kinetochore microtubule stubs might promote efficient chromosome bi-orientation through capture of pre-formed k-fibers by astral microtubules [14, 15, 17, 18] or by lateral interactions with an interpolar spindle scaffold that forms soon after nuclear envelope breakdown [75] and will give rise to bridging fibers [55] (Fig. 7E).

Our findings also reveal that Augmin impacts kinetochore microtubule turnover and poleward flux. While the role of Augmin in promoting microtubule turnover is consistent with *de novo* microtubule nucleation from pre-existing kinetochore microtubules, how Augmin promotes poleward flux remains less clear. One possibility is that Augmin-mediated microtubule amplification promotes the long-term survival of kinetochore microtubules that slide polewards, thereby facilitating tubulin incorporation at the kinetochores. Alternatively, Augmin might promote flux through its role in the formation/amplification of interpolar microtubules (as shown in the present study; see also [76]), which are critical mechanical elements necessary for poleward flux [64]. Strikingly, our live-cell super-resolution tracking of EB3-comets in the vicinity of the kinetochore allowed us to follow with unprecedented spatiotemporal resolution microtubule growth events on individual k-fibers and revealed a wide angular dispersion relative to the k-fiber axis. This is somewhat reminiscent of the “fir-tree” structure observed on k-fibers in *Haemanthus* endosperm spindles [77] and consistent with Augmin-mediated branched microtubule nucleation from pre-existing microtubules [25-28, 65, 66, 78, 79]. In agreement, Augmin depletion significantly reduced microtubule growth events within individual k-fibers. This important structural aspect has been overlooked in previous models of k-fiber formation and maturation in mammals and implies that additional factors are involved either on bundling or stabilizing newly nucleated kinetochore microtubules into a cohesive fiber. HURP has been proposed to be such a factor [48, 50], but our phenotypic analysis of k-fibers in HURP-depleted Indian muntjac fibroblasts failed to confirm this hypothesis. In addition, as opposed to critical roles of the chromosomal passenger complex, important players in the Ran-GTP pathway (e.g. TPX2) previously implicated in branched microtubule nucleation in *Xenopus* egg extracts and after *in vitro* reconstitution from purified *Xenopus* components [27, 79], appear to be largely dispensable for robust k-fiber formation and spindle assembly in Indian muntjac fibroblasts, in line with previous findings in *Drosophila* cells [8, 80]. More specifically, TPX2 has also been shown to be dispensable for microtubule branching in *Drosophila* cells [25] and after *in vitro* reconstitution from purified *Drosophila* components [66].

Curiously, astral microtubules grew much longer in the absence of Augmin and Ndc80 when compared to controls, which we interpreted as a consequence of an increased soluble pool of tubulin due to compromised k-fiber formation. Although a more direct role of Augmin and Ndc80 in centrosome-dependent microtubule nucleation cannot be formally excluded, our findings in cells contrast with previous reports that suggested that both complexes promote astral microtubule nucleation from purified centrosomes *in vitro* [33]. It was also surprising that after interference with Augmin function, PRC1 decorated parallel microtubules. This might be explained by a significant reduction of antiparallel interpolar microtubules after Augmin perturbation, which would favour PRC1 association with parallel microtubules that normally have higher off-rates (and equivalent on-rates) compared to antiparallel microtubules [81]. Indeed, recent optogenetic experiments have shown that, during recovery, PRC1 first relocates indiscriminately to the entire spindle, and only then concentrates on overlapping antiparallel microtubules [82].

Lastly, our laser-mediated partial k-fiber injury/repair assay directly demonstrates a role for Augmin in microtubule amplification from pre-existing kinetochore microtubules. Together with our measurements of kinetochore microtubule turnover and direct observation of microtubule growth events within individual k-fibers, our data provide definitive evidence for a role of Augmin-mediated microtubule amplification in k-fiber maturation, while reconciling this mechanism with growing evidence for centrosome-independent kinetochore-mediated microtubule formation during spindle assembly in animal cells, including human [14, 16]. This underscores the added value of Indian muntjac cells as a powerful model system to study mitosis, offering an important and open access resource for the cell division community.

## Acknowledgments

We would like to thank all colleagues that kindly shared invaluable reagents used in this study, Bárbara Amorim and Marbélia Fernandes from the Information Systems and Technology Unit at i3S for help with the screening website, Naoyuki Okada for guidance regarding centrinone treatment in Indian muntjac fibroblasts, and Maiato Lab members for suggestions throughout this project and the critical reading of the manuscript. Work in the Maiato lab is funded by the European Research Council (ERC) consolidator grant CODECHECK, under the European Union’s Horizon 2020 research and innovation programme (grant agreement 681443), Fundação para a Ciência e a Tecnologia of Portugal (PTDC/MED-ONC/3479/2020), and the NORTE-01-0145-FEDER-000051 project supported by NORTE 2020 under the PORTUGAL 2020 Partnership Agreement through the European Regional Development Fund.

## Author contributions

Methodology (AJP, ACA, JO, DD, LPC, PA, JD, HAL, DML); Investigation, Formal Analysis and Validation (ACA, JO, AJP, PA); Visualization (ACA, JO, AJP, HM); Writing – Original Draft (ACA, HM); Writing – Review and Editing (ACA, AJP, JO, HM); Conceptualization, Supervision, Project Administration and Funding acquisition (HM).

## Supplemental Figure Legends

**Fig. S1**. (relative to Fig. 1) **Mitotic timings of Indian muntjac fibroblasts upon gene-specific RNAi-mediated depletion.** Indian muntjac fibroblasts stably expressing H2B-GFP and labeled with 50 nM SiR-tubulin were acquired by confocal spinning-disk microscopy every 2 minutes. Mitotic duration was determined by measuring the time between NEBD to AO; shown in min. Each data point corresponds to one cell; mean ± S.D.; *p≤0.05, **p≤0.01, ***p≤0.001, ****p≤0.0001, no asterisk corresponds to not significantly different from controls.

**Fig. S2**. (relative to Fig. 2) **Augmin recruits γ-tubulin to the spindle region and is required for the formation of robust k-fibers in Indian muntjac fibroblasts.** Immunofluorescence images of control and HAUS6-depleted Indian muntjac fibroblasts labelled for α-tubulin (green, **A**), γ-tubulin (green, **B**), Mad2 (green, **C**), HURP (magenta, **D**), anti-centromere antiserum (ACA, magenta, **A-C**) and DAPI (white). Spindle length was calculated in **A’** (n= 37 Control cells; n= 28 siHAUS6 cells); overall γ-tubulin fluorescence intensity was measured and normalized to the average control levels in **B’** (n= 36 Control cells; n= 24 siHAUS6 cells); ratio between Mad2 positive kinetochores and ACA fluorescence values was determined in **C’** (n= 252 Control kinetochores; n= 176 siHAUS6 kinetochores) and HURP signal in the spindle of HAUS6- and Ndc80-depleted cells was normalized to the average control levels in **D’** (n= 65 Control cells; n= 67 siHAUS6 cells; n= 57 siNdc80 cells). **E,** cold treatment at 4°C for 5 minutes was performed to selectively destabilize non-kinetochore microtubules. α-tubulin (green) signal was normalized to the average control levels in **E’** (n= 54 Control cells; n= 31 siHAUS6 cells; n= 17 siNdc80 cells). Depletion of Ndc80 was used as a positive control (see Fig. S3C, D). The box plot determines the interquartile range; the line inside the box represents the median. **p≤0.01, *** p≤0.001, ****p≤0.0001. Scale bars: 5 µm.

**Fig. S3**. (relative to Fig. 2) **Depletion of a different Augmin subunit (HAUS1) resulted in short spindles and loss of γ-tubulin recruitment to the spindle region**. **A,** representative CH-STED image of HAUS6 localization in Indian muntjac mitotic spindle. DAPI (inverted grayscale), α-tubulin (magenta), HAUS6 (green). **B,** immunofluorescence of HAUS1-depleted cells. DAPI (white), α-tubulin/γ-tubulin (green) and ACA (magenta). Ndc80-depleted cell incubated with MG-132 for 1.5 hours and stained with anti-HURP (magenta), anti-α-tubulin (green) and DAPI (white) is shown in **C. D,** a cold treatment at 4°C for 5 min was performed to selectively destabilize non-kinetochore microtubules. α-tubulin (green), ACA (magenta) and DAPI (white). For quantifications see Fig. S2C’, D’. Scale bars: 5 µm.

**Fig. S4**. (relative to Fig. 3) **Perturbation of k-fiber formation in Indian muntjac cells is sufficient to bias tubulin polymerization towards astral microtubules. A,** representative CH-STED images of control, HAUS6-, Ndc80-, chTOG- and CLASP1/2-depleted cells incubated with 1 µM nocodazole for 2 hours, washed-out into warm medium and fixed after 2, 5 and 10 minutes. Cells were then stained for α-tubulin (inverted grayscale). Astral microtubule length was determined in **B** (Control: 2’ n= 567, 5’ n= 650, 10’ n= 434 astral microtubules; siHAUS6: 2’= 376, 5’ n= 633, 10’ n= 754 astral microtubules; siNdc80: 2’ n= 425, 5’ n= 704; 10’ n= 469 astral microtubules; siChTOG: 2’ n= 261, 5’ n= 194, 10’ n= 217 astral microtubules; siCLASP1/2: n= 161, 5’ n= 183, 10’ n= 284 astral microtubules). Acquisition ROI for siCLASP1/2 nocodazole washout 2’ image is represented in grey-dashed line. Mean ± S.D.; ns: not significant, **p≤0.01, ****p≤0.0001 relative to controls. Scale bar: 5 µm.

**Fig. S5**. (relative to Fig. 4) **Ndc80 depletion does not affect microtubule re-growth in acentrosomal spindles. A,** representative CH-STED image of an Indian muntjac cell stably expressing GFP-Centrin-1 (magenta) and 2xGFP-CENPA (magenta), treated with 1 µm nocodazole for 2 hours. No microtubules were detected (α-tubulin, green). **B,** representative CH-STED image of an Indian muntjac cell stably expressing GFP-Centrin-1 (magenta) and 2xGFP-CENPA (magenta), 2 minutes after washout from nocodazole, upon 8-days DMSO treatment. Yellow arrowheads indicate the centrioles. **C,** microtubule regrowth assay after 2, 5 and 10 minutes nocodazole washout in warm medium, upon 8-days centrinone treatment and Ndc80 RNAi transfection for 24 hours. Cells were fixed and immunostained with α-tubulin antibody (green) and DAPI (inverted grayscale). Insets show 2.5X magnification of selected regions with kinetochore and nucleated microtubules (grayscale for single channels of CENP-A and α-tubulin). The percentage of kinetochores with microtubules, **D**, and overall microtubule length, **E**, at each time point is shown (siNdc80 2’ n= 12 cells/112 microtubules; 5’ n= 21 cells/280 microtubules; 10’ n= 14 cells/142 microtubules). Each data point represents one cell (**D**) or one microtubule (**E**). Mean ± S.D.; ns: not significant, *p≤0.05, ****p≤0.0001. Scale bars: 5 µm.

**Fig. S6**. (relative to Fig. 6) **K-fibers in TPX2-, HURP-, chTOG and CLASP1/2-depleted cells were largely indistinguishable from controls. A,** comparison of confocal and CH-STED live images of EB3-Halotag/JF646. Full width at half maximum (FWHM) was used as a proxy of spatial resolution: confocal ∼300 nm; CH-STED ∼100nm. **B,** representative super-resolution images of TPX2-, HURP-, chTOG- and CLASP1/2-depleted cells stained with α-tubulin and DAPI (white; opacity 15%). Temporal color code tool on Fiji was used to correspond each α-tubulin z-plane to a different color. Scale bar: 5 µm. Insets show the max-projection (selected z-planes) of a k-fiber (α-tubulin, inverted grayscale). Scale bar: 1 µm. **C,** live CH-STED imaging conditions did not result in any obvious phototoxicity or relevant photobleaching throughout the recordings confirmed by the comparison of pre- and post-recording images of control, siHAUS6 and siNdc80 treated cells. EB3-Halo tag/JF646 (green), GFP-CENP-A (magenta). Scale bar: 5 µm. **D,** EB3 comets velocity was determined from the kymographs shown in Fig. 6C. ns: not significant. **E,** cross-correlation analysis of EB3 comets in control cells. Analyzed region of the k-fiber is shown on the right (yellow box). Raw (left) and 2-sigma filtered (right) next-neighbor cross-correlation function averaged over time-frame pairs for the ROI shown of the left. Flow angle and average velocity were determined based on the center of mass of the filtered correlation spot. The polar graph (right) shows the semi-infinite domain Radon transform used to estimate the trajectories’ angular dispersion (50% level).

**Fig. S7**. (relative to Fig. 7) **Validation of the laser microsurgery-based k-fiber ablation assay**. **A,** representative spinning-disk confocal images of total k-fiber severing in control Indian muntjac fibroblasts stably expressing GFP-α-tubulin (green) and mScarlet-CENP-A (magenta). This resulted in two fully separated kinetochore- and pole-proximal microtubule stubs that showed independent pivoting movements, followed by the rapid poleward movement of the kinetochore-proximal stub after being captured by neighboring microtubules Pre-cut frame is shown on the left. Yellow dashed-rectangle indicate the ablated k-fiber. Insets show the analyzed k-fibers (GFP-α-tubulin, inverted grayscale). Scale bar: 5 µm. -10 sec represents arbitrary time before partial k-fiber ablation. Purple arrowhead points to the ablated k-fiber portion at time zero (first frame after laser ablation). Scale bar: 1 µm, time: min:sec. **B,** immunoblot analysis of lysates from Indian muntjac cells stably expressing α-tubulin-GFP/mScarlet-CENP-A treated with control or partial HAUS6 RNAi. An antibody against HAUS6 was used to validate siRNA mediated knockdown (upper band, ∼26% depletion; asterisk indicates the band of interest). α-tubulin was used as loading control. **C,** partial k-fiber damage was confirmed by correlative live-cell spinning disk confocal microscopy (left) and CH-STED nanoscopy (right) upon fixation and immunostaining of the damaged cell. α-tubulin-GFP (green), mScarlet-CENP-A (magenta). -10 sec corresponds to arbitrary time before the k-fiber ablation (Pre-Cut). 0 sec corresponds to the first frame after k-fiber ablation. Yellow arrowhead points towards the ablated region. After laser-mediated k-fiber injury, the same cell was fixed and stained with α-tubulin (inverted grayscale) and DAPI (inverted grayscale). Scale bar: 5 µm.

## Legends for Supplemental Movies S1 to S3

**Movie S1. Augmin is required for k-fiber formation and chromosome dynamics.** CH-STED time-lapse imaging of control, HAUS6- and Ndc80-depleted Indian muntjac fibroblasts stably expressing GFP-CENP-A (magenta) and EB3-Halotag, conjugated with a far-red dye JF646 (green). Cells were treated for 30 minutes with MG-132 and imaged every 8 seconds (sec) for two minutes. Scale bar: 1 µm.

**Movie S2. Augmin is essential for microtubule growth within k-fibers.** Live CH-STED time-lapses of control, HAUS6- and Ndc80-depleted Indian muntjac fibroblasts stably expressing GFP-CENP-A (magenta) and EB3-Halotag, conjugated with a far-red dye JF646 (green), show microtubule polymerization events within one k-fiber. Time-lapse was 750 ms. Time is displayed in seconds (sec). Scale bar: 1 µm.

**Movie S3. Augmin promotes k-fiber repair after a partial k-fiber injury.** Spinning-disk confocal movies of control and partial HAUS6-depleted Indian muntjac fibroblasts stably expressing GFP-α-tubulin (green) and mScarlet-CENP-A (magenta), illustrating k-fiber recovery after partial k-fiber laser ablation. Whole-spindle image sequences were stabilized for any movement using the spindle poles as a reference. Cells were imaged every 1 second. Time is displayed in minutes:seconds -10 seconds corresponds to arbitrary time before laser k-fiber injury. White arrow points to the ablated k-fiber portion. Scale bar: 5 µm.

## Materials and Methods

### Molecular Biology

pLVX-EB3-Halotag, pRRL-2xGFP-CENPA, pLVX-mScarlet-CENPA and pLVX-PAGFP-tubulin were generated by Gibson assembly. pLVX-GFP-Centrin-1 was a gift from Manuel Thery (Addgene #73331).

### Cell Culture and Lentiviral Transduction

All Indian muntjac cell lines were grown in Minimum Essential Media (MEM) (Corning), supplemented with 10% FBS (GIBCO, Life Technologies) at 37°C in humidified conditions with 5% CO_2_. Indian muntjac hTERT-immortalized fibroblasts were a gift from Jerry W. Shay [43] and Indian muntjac stably expressing H2B-GFP were previously generated in [38]. For microtubule plus-ends, centrioles, CENP-A, photoactivatable tubulin and α-tubulin tagging, cells were transduced with pLVx-EB3-Halotag, pLVx-GFP-Centrin-1, pRRL-2xGFP-CENPA, pLVx-mScarlet-CENPA, pLVx-pAGFP-tubulin and pRRL-EGFP-α-tubulin [83] lentiviral plasmids, respectively. Lentivirus particles were added to the standard culture media with 1:2000 Polybrene (Sigma) for 12h. Stable lines with uniform level of expression and sufficient fluorescence intensity were selected by FACS. To select pLVx-EB3-Halo tag positive cells, 20 nM of the far-red dye JF646 (Promega®) was added 5-10 minutes before FACS sorting.

### Identification of Indian Muntjac Sequences

The protein sequences of human genes were obtained from NCBI and used as query for tblastn (version 2.2.29 [84]) using the Indian muntjac genome scaffold sequences and predicted coding sequences (CDC) as targets. Sequence alignments with at least 80% identity, highest coverage of human genes, with matching scaffold and CDS intervals from both tblastn runs were used to identify Indian muntjac orthologs of human gene sequences.

### Design of siRNAs for RNA Interference (RNAi)

The design of the siRNA sequences was performed using the application BLOCK-ITTM RNAi Designer (Thermo Fisher Scientific). We provided the nucleotide sequence of the genes of interest, selected an ideal CG percentage between 35%–55% and the recommended default motif pattern for the RNAi design. From the 10 designs generated, the one with higher probability of knockdown was selected for each protein of interest.

### siRNA Experiments

For siRNA experiments, Indian muntjac fibroblasts were plated at 60%–70% confluence in 6-well plates or 22x22 mm no. 1.5 glass coverslips (Corning), previously coated with fibronectin (Sigma-Aldrich) as described in [85] for 24 hours in normal medium. Cells were then starved with MEM supplemented with 5% FBS for 30 minutes. siRNA transfection was performed using 5 μL of Lipofectamine RNAi Max (Invitrogen) and 50-100 nM of the respective siRNA, each diluted in 250 μL of serum free-medium (Opti-MEM, Gibco). Untreated and mock transfection (with lipofectamine only) were indistinguishable and therefore referred to as “Control”. Cells were analyzed 24, 48, 72 or 96 hours after depletion, depending on the protein of interest. Depletion efficiency was monitored by western blotting and phenotypic analysis. 65 RNAi sequences were optimized and are available in http://indianmuntjac.i3s.up.pt. 66 conditions were analysed in this study (including double depletion of CLASP1/2 and VASH1/2). Cyclin-B depletion prevented cells’ entrance in mitosis, so it was not included in the phenotypical/clustering analyses of this study.

### Western Blotting

Indian muntjac fibroblasts were collected by scraping the adherent cells or by trypsinization and centrifuged at 200 x g for 5 minutes. The pellet was washed with PBS and centrifuged again. Cells were then resuspended in lysis buffer (*NP-40:* 20 nM HEPES/KOH,pH 7.9; 1 mM EDTA,pH 8; 1 mM EGTA; 150 mM NaCl; 0.5% NP40-IGEPAL; 10% glycerol; 2 mM DTT, supplemented with 1:50 protease inhibitor and 1:100 phenylmethylsulfonyl fluoride (PMSF); OR *for DNA-binding proteins*: 50mM Tris-HCl, pH 7.4; 0,1% digitonin (in EtOH); 0.5% Triton; 400 nM NaCl, supplemented with 30 µg/mL RNAse, 20 µg/mL DNAse, 10 µM MgCl_2,_ with 1:50 protease inhibitor and 1:100 PMSF). The samples were snap-frozen in liquid nitrogen and kept on ice for 30 minutes. After centrifugation at 20 800 x g for 15 minutes at 4°C, protein concentration was determined by Bradford protein assay (Thermo Fisher Scientific). Protein lysates were run on 7.5/10/15% SDS-PAGE (25-50 µg/lane) according to their molecular weight and transferred to a nitrocellulose Hybond-C membrane using an iBlot Gel Transfer Device (Bio-Rad) or using a wet-transfer system (if protein molecular weight >120kDa). Membranes were then blocked in 5% milk diluted in PBS 0.05% Tween and the primary antibodies were incubated overnight at 4°C at the dilutions shown in Table S1. After successive washes, the membrane was incubated with the secondary antibodies for 1 hour at RT - 1:5000 α-mouse-HRF; α-rabbit-HRF; α-sheep-HRP (Jackson ImmunoResearch). Detection was performed with Clarity Western ECL Substrate (Bio-Rad). Quantification of blots was performed with a Bio-Rad ChemiDoc XRS system using the IMAGE LAB software and immunosignals were normalized to GAPDH, α-tubulin or vinculin expression depending on protein molecular weight.

### Immunofluorescence

Indian muntjac fibroblasts were seeded on fibronectin coverslips 24h before the experiment, as shown in [85]. Cell were fixed with ice-cold methanol (Sigma) for 4 minutes at -20°C; or 4% paraformaldehyde (Electron Microscopy Sciences) or, for STED microscopy, PFA 4% supplemented with 0.1%-0.2% glutaraldehyde (Electron Microscopy Sciences) for 10 minutes at room temperature (RT). Autofluorescence was quenched by a 0.1% sodium borohydride solution (Sigma-Aldrich) after aldehyde fixation. Extraction after paraformaldehyde fixation was performed using PBS-0.5%Triton (Sigma-Aldrich) for 10 minutes. Cells were incubated with blocking solution (10% FBS diluted in PBS-0.05%Tween 20) or in cytoskeleton buffer (274 mM NaCl, 10 mM KCl, 2.2 mM Na_2_HPO_4_, 0.8 mM KH_2_PO_4_, 4 mM EGTA, 4 mM MgCl_2_, 10 mM Pipes, 10 mM glucose, pH 6.1; with 0.05%Tween 20) for 1 hour at RT, followed by incubation with primary antibodies in the same solution over-night at 4°C (for antibodies’ dilutions check Table S1). Subsequently, cells were washed 3x with PBS-0.05% Tween and incubated for 1 hour at RT with the corresponding secondary antibody - Alexa 488, 568 and 647 (Invitrogen); or Abberior STAR 580 and Abberior STAR (Abberior Instruments) for STED microscopy. For STED microscopy, both primary and secondary antibodies were used at 1:100-150 concentrations. After adding 1 µg/mL 4’6’-diamidino-2-phenylindole (DAPI) in PBS-0.05% Tween (Sigma Aldrich) for 5 minutes, coverslips were washed in PBS and sealed on glass slides mounted with 20 mM Tris pH8, 0.5 N-propyl gallate, 90% glycerol.

### Drug Treatments

Mitotic arrest at metaphase was obtained using 3-5 µM MG132 (Calbiochem). Live-cell and fixed cell analysis using MG132 was performed in the first 2 hours after drug addition to avoid cohesion fatigue. Microtubule depolymerization was triggered using 1 µM of nocodazole for 2 hours before fixation or washout. Microtubule re-growth assay was performed by washing out nocodazole with warm medium 2, 5 and 10 minutes before fixation. To induce centriole loss due to Plk4 inhibition, an 8-days treatment with 125 nM centrinone was performed. An equivalent volume of DMSO was used as control for each drug treatment. For the live-CH-STED experiments 75 nM JF646 was conjugated with Halo-tag expressing Indian muntjac fibroblasts. SiR-tubulin and SiR-DNA (Spirochrome) [44] were used to visualize microtubules and chromosomes, respectively, at 50 nM concentration incubated for 1 hour prior to live-cell imaging.

### Time-lapse Spinning-disk Confocal Microscopy

Indian muntjac fibroblasts stably expressing human H2B-GFP were plated on fibronectin coated 22x22 mm no. 1.5 glass coverslips 24h before imaging. 1 hour before live-cell imaging, cells were incubated in Leibovitz’s L15 medium (GIBCO, Life Technologies) with SiR-tubulin cell-permeable dye. Coverslips were assembled onto 1-well Chamlide CMS imaging chambers (Microsystem AB; Sweden) immediately before imaging. Live-cell imaging was performed on a temperature-controlled Nikon TE2000 microscope equipped at the camera port with a Yokogawa CSU-X1 spinning-disc head (Solamere Technology), an FW-1000 filter-wheel (ASI) and an iXon+ DU-897 EM-CCD (Andor). The excitation optics are composed of two sapphire lasers at 488 nm and 647 nm (Coherent), which are shuttered by an acousto-optic tunable filter (Gooche&Housego, model R64040-150) and injected into the Yokogawa head via a polarization-maintaining single-mode optical fiber (OZ optics). Sample position was controlled by a motorized SCAN-IM stage (Marzhauser) and a 541.ZSL piezo (Physik Instrumente). The objective was an oil-immersion 60x 1.4 NA Plan-Apo DIC CFI (Nikon, VC series), yielding a 190 nm/pixel sampling. All image acquisition was controlled by NIS Elements AR software. An image stack (9 planes separated by 1.5 µm) was acquired every 2 minutes, spanning a total depth of 12 µm.

### Microtubule Turnover Measurements by Photoactivation

Microtubule turnover was measured in Indian muntjac fibroblasts stably expressing pAGFP-Tubulin/GFP-Centrin-1, seeded on fibronectin coated 22x22 mm no. 1.5 glass coverslips. Medium was changed to Leibovitz’s L15 medium with SiR-DNA cell-permeable dye (Spirochrome) 1 hour before live-cell imaging. Mitotic cells were identified by Differential Interference Contrast (DIC) microscopy and imaging was performed using a Plan-Apo 100× NA 1.40 DIC objective on a Nikon TE2000U inverted microscope equipped with a Yokogawa CSU-X1 spinning-disc confocal head containing two laser lines (488 nm and 647 nm) and a Mosaic (Andor) photoactivation system (405 nm). Photoactivation was performed in cells with all chromosomes aligned at spindle equator, identified by SiR-DNA signal. Microtubules were locally activated on a thin stripe of ∼1 μm width spanning one half-spindle in an area mid-way between the spindle pole and the chromosomes. The 405 nm laser was used at 75% power and cells were pulsed once (500 ms exposure). Seven 1-μm fluorescence image planes were captured using a 100X oil-immersion 1.4 numerical aperture objective every 15 seconds for 4.5 minutes. To determine fluorescence dissipation after photoactivation (FDAPA), whole-spindle sum-projected kymographs were generated and quantified using a custom-written MATLAB script. Intensities were normalized to the first time-point after photoactivation (following background subtraction from the respective non-activated half-spindle). Values were corrected for photobleaching by normalizing to the fluorescence loss of whole cell sum projected images. To calculate microtubule turnover, the normalized intensity values at each time point were fitted to a double exponential curve *A*1 × *e*^−*k*1×*t*^ + *A*2 × *e*^−*k*2×*t*^; t - time, A1 - less stable microtubule population (non-kMTs); A2 - more stable microtubule population (kMTs); k1 and k2 - decay rates of population fractions A1 and A2, respectively (only fittings with R^2^>0.98 were retained). From these curves, the rate constants, and the percentage of microtubules for the fast - typically interpreted as the fraction corresponding to non-kMTs- and the slow - typically interpreted as the fraction corresponding kMTs - processes were obtained. The half-life time was calculated as ln(2)/k for each microtubule population.

### Flux Velocity Measurements

To determine microtubule poleward flux velocity, the whole-spindle sum-projected image sequence was first stabilized using the spindle centrosomes (the coordinates of which were previously determined using a simple centroid-based tracking routine) as references. This procedure generates a guided-kymograph, where a virtual spindle equator remains static (i.e., without translation or rotation) throughout time. The distance between the photoactivated stripe and the virtual equator, as determined by the midpoint between centrosomes, yields the poleward flux velocity.

### K-fiber Maturation Assay – Laser Microsurgery

Indian muntjac fibroblasts stably expressing human EGFP-α-tubulin and mScarlet-CENP-A were plated on fibronectin coated Ø 25 mm no. 1.5 glass coverslips. Before live-cell imaging, cells were incubated in Leibovitz’s L15 medium (GIBCO, Life Technologies). Laser microsurgery was performed on an inverted microscope (TE2000U; Nikon) with a doubled-frequency laser (FQ-500-532; Elforlight), focused by a 100× 1.4 NA plan-apochromatic DIC objective lens (Nikon) equipped with an iXonEM+ EM-CCD camera (Andor Technology). One plane was acquired every 1 second for 2 minutes and subsequently every 1 minute up to 5 minutes. Partial disruption of k-fibers was performed by 2-5 consecutive pulses (0.35 μm step between pulses) conjugated with 3 pulses (0.4 μm Z-step) at each point (12 Hz repetition rate). The pulse width was 10 ns and the pulse energy was 1.5–2 μJ. For the quantification, a straight-line with a specific width (consistent with k-fiber width) was outlined in the ablated k-fiber and monitored during the first 30-40 seconds of movie. All data was normalized to the k-fiber intensity values pre-cut at each time point. The intensity line profiles were then analyzed, and two different regions were defined: 1) stub – corresponding to the region between the kinetochore and the point immediately before the sharp intensity drop; 2) dip – corresponding to a 450 nm region following the stub which contains the ablated k-fiber portion. A region of 300 nm was used as a spacer between the stub and the dip to exclude the contribution of microtubule flux (see scheme Fig. 7A). To accommodate the focal-plane fluctuations, the average dip intensity was normalized to the average stub intensity at each time point. Fluorescence Recovery After k-fiber Severing (FRAS) was determined by fitting the ratio at each point to a one phase decay (least squares fit): *y* = (*y*0 − *plateau*) × *e*^−*kt*^ + *plateau*); *y0*= intensity at time zero; *K*= ln(2)/half-life; *t*= time (seconds). Healing recovery percentage was calculated using the following equation: R= ((I_tf_ -I_ti_)/(1 -I*ti*)) ×100, where R= healing recovery; *I=* dip/stub intensity ratio; *tf*= for controls, corresponds to the time were the ratio equals 1; for HAUS6, corresponds to time 30 seconds (4 seconds after all control treated cells reached a ratio of 1); *ti*= time zero/post-cut. The analysis was restricted to cells where stub intensity variation<20% and dip intensity drop>15%. Only a minor fraction of the analyzed k-fibers belonged to the big kinetochore as it rarely localizes on the periphery of the spindle (ideal place to perform k-fiber surgery and track the outcome) and easily gets out of focus (due to its large size). For correlative CH-STED nanoscopy after microsurgery coverslips were assembled onto perfusion imaging chambers (Ske, Research Equipment) before imaging. Immediately after microsurgery, cells were perfused with 5 mL of 4% PFA + 0.2% glutaraldehyde in CBS and stained as described in methods section “*Immunofluorescence”.* The surgery laser was used to mark a reference frame in the coverslip glass after fixation, thus allowing the cell of interest to be located using a 10x objective on the STED microscope.

### Stimulated Emission Depletion (STED) Microscopy and Quantification

For Coherent-Hybrid STED (CH-STED) imaging, an Abberior ’Expert Line’ gated-STED microscope was used, equipped with a Nikon Lambda Plan-Apo 1.4NA 60x objective lens. CH-STED was implemented as described before [42]. All acquisition channels (confocal and CH-STED) were performed using a 0.8 Airy unit pinhole. A time-gate threshold of 500 ps was applied to the STED channel to avoid residual confocal-resolution signal contribution. Fixed-cell images were acquired using excitation wavelengths at 561 nm and 640 nm. Excited volumes were doughnut-depleted with a single laser at 775 nm. Pixel size was set to 40 nm. All images show max-intensity projections. For representation purposes, in Fig. 6A and Fig. S6B, a temporal color code tool in Fiji (ImageJ) was used to correspond each z-plane to a different color and DAPI channel was added in Adobe Photoshop CS6 as a separate layer with an opacity of ∼15%. Astral microtubule length was measured as the curve length between the spindle pole and the microtubule distal tip in maximum-projection images, using the segmented line tool in Fiji. Microtubule length and number after nocodazole washout in centrinone-treated cells was determined by measuring the distance from a kinetochore to the distal microtubule tip and by counting the number of detected microtubules foci per kinetochore, respectively. To determine the proportion of kinetochore-attached microtubules fraction in the spindle, two volumes that are assumed to contain well-defined microtubule populations were defined. *Region 1*, the non-kinetochore microtubules source, was defined as the volume between two surfaces, each one generated by interpolation of the array of kinetochores in either half-spindle (using a “thin-plate” spline interpolant). The kinetochores were previously defined manually through identification of microtubule-fiber ends in Fiji. To quantify Region 1 signal, instead of using the whole volume between the two surfaces, we chose a sub-plate of 200 nm width. Eventual microtubules crossing this plate that go on to attach to a kinetochore are assumed to be very rare, particularly in late prometaphase and metaphase. We then assumed that the microtubules accounted for in Region 1 generally extend outside it, even if slightly. An equivalent assumption is that the interpolar microtubules density does not change significantly between Region 1 and its immediate (<500 nm) vicinity. Accordingly, we defined a Region 2 volume lying poleward relative to kinetochores, which contains the sum of two microtubule populations (those that couple the half-spindles and those that attach to kinetochores). Specifically, it is defined as 200 nm-wide and 100 nm poleward-shifted replica of the surfaces defined above (for both half-spindle kinetochores), which enclosed Region 1. The final result for the fraction *f* of kinetochore-attached microtubules is f=1-S_Region1_/S_Region2_, where S is the background-subtracted integrated photon count in the corresponding region. Background level was estimated as the average photon count in a metaphase plate region that was visually identified as being devoid of microtubule signal. Total tubulin intensity in HAUS6-depleted cells was normalized to the average levels of control cells. To visualize live microtubule growing events within a single k-fiber, we used Indian muntjac fibroblasts stably expressing EB3-Halotag/GFP-CENPA imaged by live-CH-STED microscopy with a 1.4NA 60x objective warm-up to 37.5 (set-point), every 750 ms or 8 seconds (to evaluate kinetochore dynamics with or without Augmin and Ndc80). Confocal images of CENP-A were acquired using an excitation wavelength of 488 nm and CH-STED images of EB3 using an excitation wavelength of 640 nm, doughnut-depleted with a single laser of 775 nm. To standardize the quantifications, only chromosomes with a large kinetochore were considered for the analysis. Pixel size was set to 40 nm. Inter-kinetochore distances were measured using a custom program written in MATLAB, which determines the distance between the vertices of parabolic fits performed on the peaks observed in 3.5 µm length line profiles averaged over a 200 nm width outlined across the pair of KTs in Fiji. To prepare the data for local kymograph analysis, we used a custom routine written in MATLAB to compensate for kinetochore movements described by the following steps: (1) 2D tracking of kinetochore tips, (2) region-of-interest (ROI) dimensions definition, and (3) thick-kymograph generation with the ROI being automatically translated and rotated every frame according to the 2D coordinates of the two reference objects in step 1. The direct output is a whole-k-fiber kymograph that can be used to generate a set of aligned images that are the basis for all subsequent analysis. A two-color guided kymograph was represented to facilitate the visualization of EB3 comets that reached the kinetochore (classified as k-fiber growing events) or EB3 comets that surpass the kinetochore (named as interpolar microtubules’ growing events). By inspection of the kymograph, a first estimate is obtained by manual definition of a kymograph stripe (in x-t), the slope of which is the translation velocity vector projected onto the spindle axis. To control for the sub-estimation of velocity incurred by the said projection, manual estimation of microtubule inclination relative to the spindle axis was used to warrant exclusion of trajectory angles higher than 25°, yielding a real velocity less than 10% higher than calculated. A refined result was obtained by running a custom MATLAB routine where an intensity-based centroid is determined at each timepoint in a preset x-neighborhood from the manual estimate. A linear fit is finally made to the collection of centroids, the slope of which yields a velocity relative to the virtual spindle equator. Chromosome poleward and anti-poleward velocity relative to the equator were measured after each EB3-Halotag track obtained from guided-kymographs aligned to the spindle pole. Velocity was determined measuring the slope of the linear movement within a half spindle. Chromosome oscillations’ amplitude was measured by subtracting the distance to the pole in the starting moment of anti-poleward to poleward movement. The related periodicity was calculated extracting the time from the start of polymerization cycle until the begging of the next polymerization cycle. The distributions shown in the scattered plots represent each track measured in the total number of cells. Finally, EB3 bursts at kinetochore were calculated dividing the number of time frames with EB3-signal at kinetochores by the total time frames in which a kinetochore was detected. In control cells, the number of EB3 bursts at kinetochores was 0.5, corresponding to the polymerization events during 2 minutes movies. To determine angular dispersion of EB3 comets within a k-fiber we used a custom Matlab script in which we outlined a ROI with approximately the width of a K-fiber, ∼500 nm from the kinetochore during the 1 minute movies. Correlation maps were constructed as an average of the individual spatial cross-correlation functions between time-neighbors (750ms time-lapse) with the original images masked for photon counts below 1 sigma. A 25 μm/minute velocity was preset as a maximal shift. The average correlation map was cleaned at a 2 sigma above average level, which was interpreted as the correlation peak used for further quantification. The peak’s center of mass yielded an average flow velocity and direction. To estimate an angular spread attributable to the different orientations of the EB3 comets trajectories, we integrated the correlation map values along radial lines (using a semi-infinite domain version of the Radon transform). The result is a comet-like polar diagram, the angular spread of which was determined as the angle separating the 50% level-crossings around the comet maximum. Finally, angular cropping of the correlation map using the 50% level limits yields the adjusted flow velocity.

### Wide-field Image Acquisition and Quantification

3D wide-field image acquisition (0.23 μm z-step) was performed on an AxioImager Z1 (60× Plan-Apochromatic oil differential interference contrast objective lens, 1.46 NA, Carl Zeiss Microimaging Inc.) equipped with a CCD camera (ORCA-R2, Hamamatsu) operated by Zen software (Carl Zeiss, Inc.). Blind deconvolution of 3D image datasets was performed using AutoquantX software (Media Cybernetics). All images show maximum intensity projections. Spindle length was determined by measuring the distance between the two spindle poles (labeled with gamma-tubulin) using the straight-line tool in Fiji. γ-tubulin intensity and α-tubulin levels after cold-treatment were determined by drawing an elliptical ROI around the spindle in sum-projected images (Fiji). Background fluorescence was measured outside the ROI and subtracted from each cell. All values were normalized to the average levels of control cells. *HURP* protein levels were determined by drawing two elliptical selections of different sizes (one containing the other) around the mitotic spindle in sum-projection images. The fluorescence intensity was background subtracted according to the equation: S_in,corrected_ = S_in_ - ((S_out_-S_in_)/(A_out_-A_in_ ))× A_in_ S : signal; A: area; and normalized to average fluorescence intensity of control cells. For quantification of cMad2 protein levels at kinetochores in metaphase arrested cells, ROI manager in Fiji was used. cMad2 fluorescence intensity was background subtracted and normalized for the levels obtained for ACA in the same kinetochore. Adobe Photoshop 2021 and Adobe Illustrator CS5 (Adobe Systems) were used for histogram adjustments and panel assembly for publication.

### Phenotypical characterization – screening analysis

In preparation for the hierarchical clustering analysis, we set the following eight binary features: A) incomplete congression and faster mitosis (if cells failed chromosome congression and mitotic duration was faster than the average NEBD-AO timing minus two standard deviations of the mean: <23 minutes ); B) incomplete chromosome and normal mitotic duration (if cells failed chromosome congression and mitotic duration was faster than the average NEBD-AO timing plus two standard deviations of the mean: <52 minutes); C) incomplete congression and prolonged mitosis (if cells didn’t congress the chromosomes and mitotic duration was slower than the average NEBD-AO timing plus two standard deviations of the mean: ≥52 minutes); D) congression delay (if cells congressed all the chromosome to spindle equator and NEBD-Metaphase time was superior to its average minus two standard deviations of the mean: ≥41 minutes); E) metaphase delay (if cells congressed all the chromosome to spindle equator and NEBD-Metaphase time was superior to its average minus two standard deviations of the mean: ≥28 minutes); F) anaphase lagging chromosomes; G) mitotic death and H) cytokinesis failure.

### Hierarchical clustering analysis

The effect of specific genes on mitosis dynamics was characterized using hierarchical clustering analysis. A set of 8 binary features was used to describe the alterations produced by silencing specific genes through siRNA. For each silenced gene, a phenotypical fingerprint was calculated as the mean value of the features vectors for all tested cells. In other words, the calculated fingerprint vector for each gene corresponds to the probabilities of occurrence of the 8 features. The phenotypes fingerprints were compared using a dendrogram representation where the Euclidean distance was used as the distance metric. Ndc80 and Spc24 were used as “positive controls” for the dendrogram cut-off definition (giving rise to 10 multi-protein clusters). This method provided an unbiased description of how the different fingerprints clustered together. All calculations and graphical representations were carried out using scripts in MATLAB 2018b (The Mathworks Inc, USA).

### Statistical analysis

Statistical analysis was performed using GraphPad Prism 8. All data represents mean ± S.D., except for Fig. S2 and Fig. 3B where median and interquartile range are shown. D’Agostinho-Pearson omnibus normality test was used to determine if the data followed a normal distribution. If α=0.05, a statistical significance of differences between the population distributions was determined by Student’s t test. If α<0.05, statistical analysis was performed using a Mann-Whitney Rank Sum test. Quantifications of kinetochore dynamics (poleward and anti-poleward velocity, chromosome oscillations’ amplitude and period) were analyzed using multiple comparisons one-way ANOVA. Quantifications of microtubule foci number and length upon centrinone/DMSO treatment were analyzed using a multiple comparison Kruskal-Wallis test. For each graph, ns= not significant, *p<0.05, **p<0.01, ***p<0.001 and ****p<0.0001, unless stated otherwise. All results presented in this manuscript were obtained from pooling data from at least 3 independent experiments except for the screening results presented for the following proteins siBubR1, siMPS1, siCdc20, siCENP-I, siCENP-F, siSurvivin, siAstrin, siEB1, siCLASP2, siCLASP1/2, siCLIP-170, siKTNB1, siAuroraA, siKif18B, siKID, siDHC, siNuMA, siNDEL1, siSpindly, siCnd1, siSgo1, siPlk1, siPRC1, siSeparase, siCDK1; quantification of γ-tubulin and α-tubulin intensity after cold treatment (Fig. S2B, B’, D, D’) and quantification of ipMTs vs kMTs proportion (Fig. 3C-E), where two independent experiments were performed.

#### Data availability

A public repository where time-lapse movies, phenotypical fingerprints and siRNA sequences for each gene can be conveniently browsed is freely available as a community resource at http://indianmuntjac.i3s.up.pt

#### Code availability

No new code was generated in this study.

