## Supplementary figures and images for "Kinetochore-mediated microtubule assembly and Augmin-dependent amplification drive k-fiber maturation in mammals"

### Figure S1

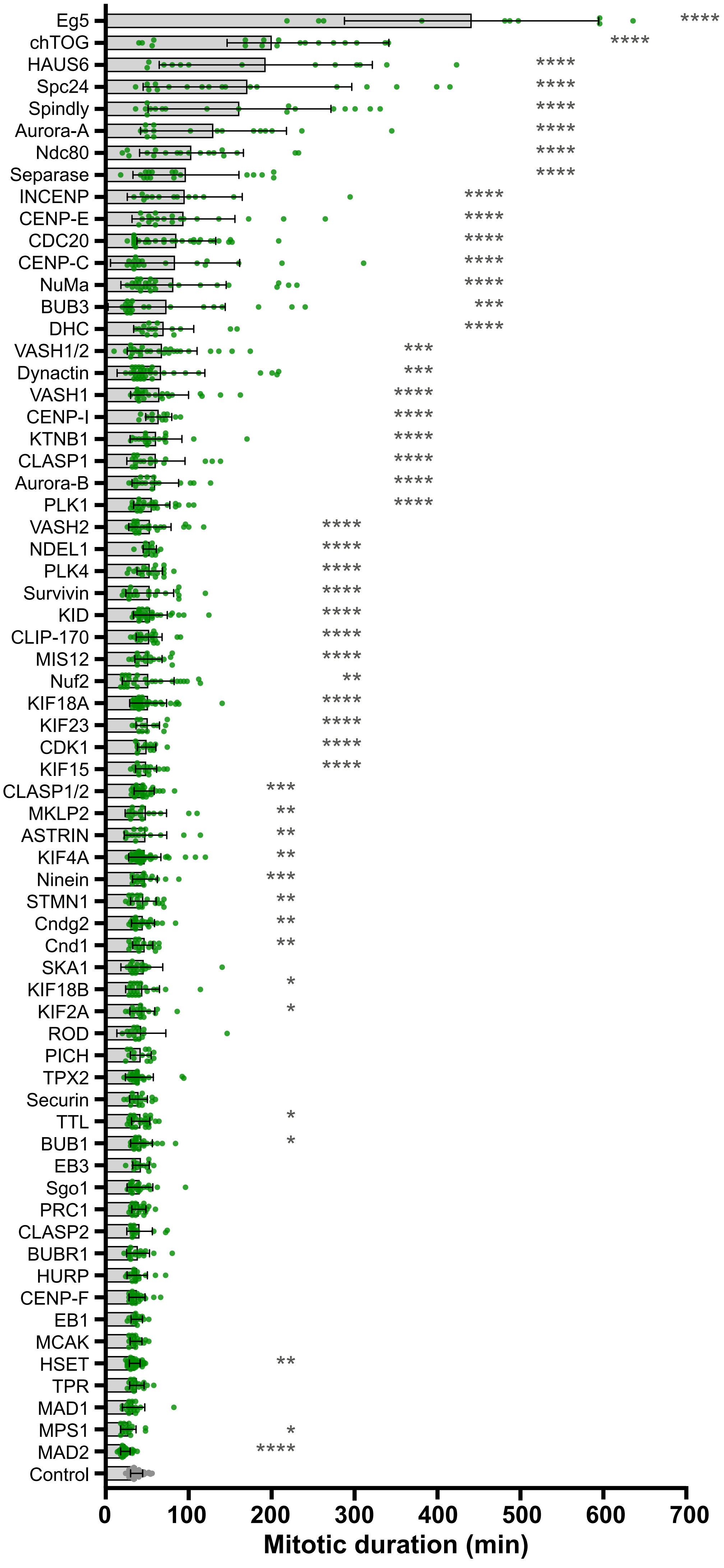


**Figure S1 - Almeida et al.**

### Figure S2

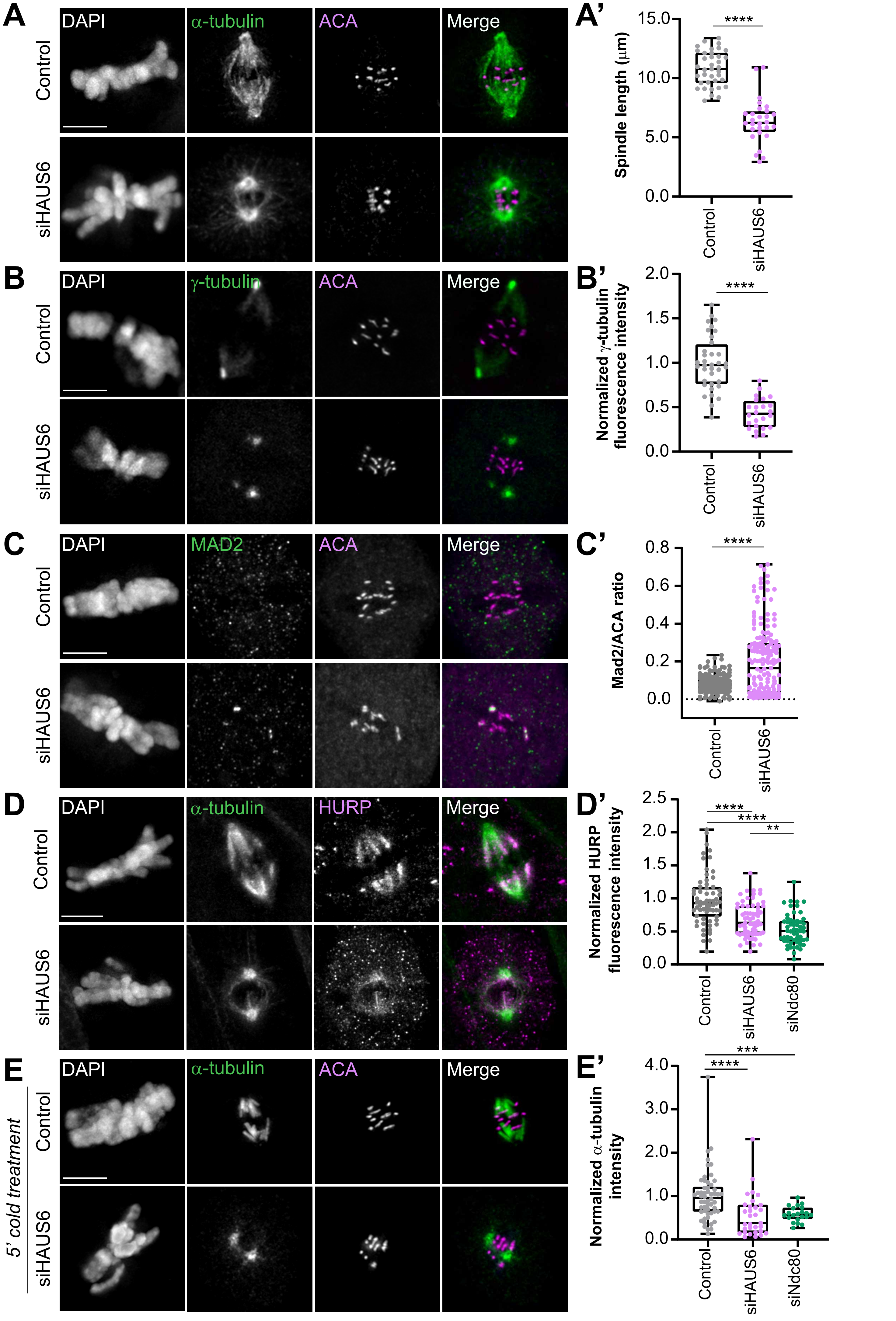


Figure S2 - Almeida et al.

### Figure S3

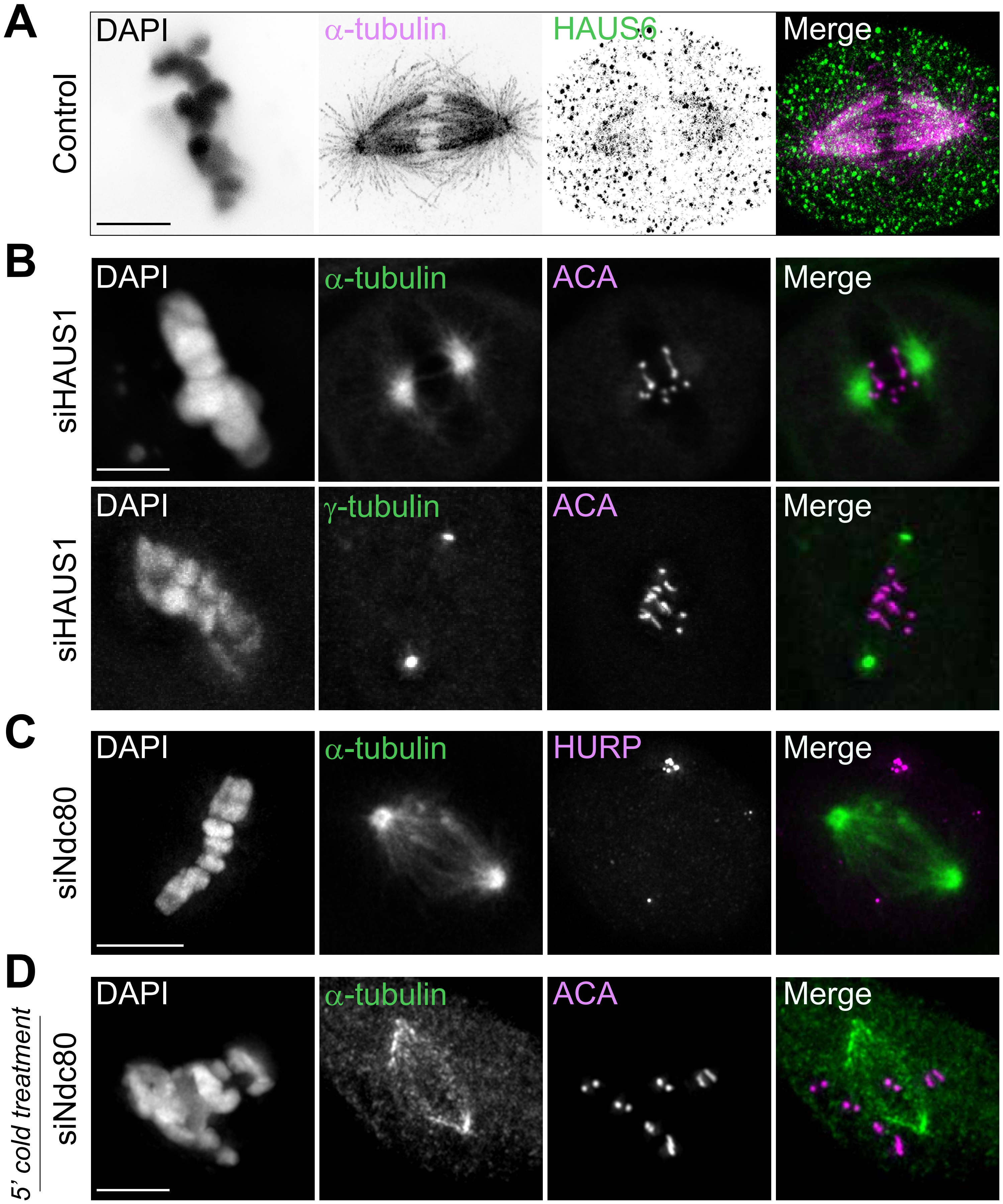


**Figure S3- Almeida et al.**

### Figure S4

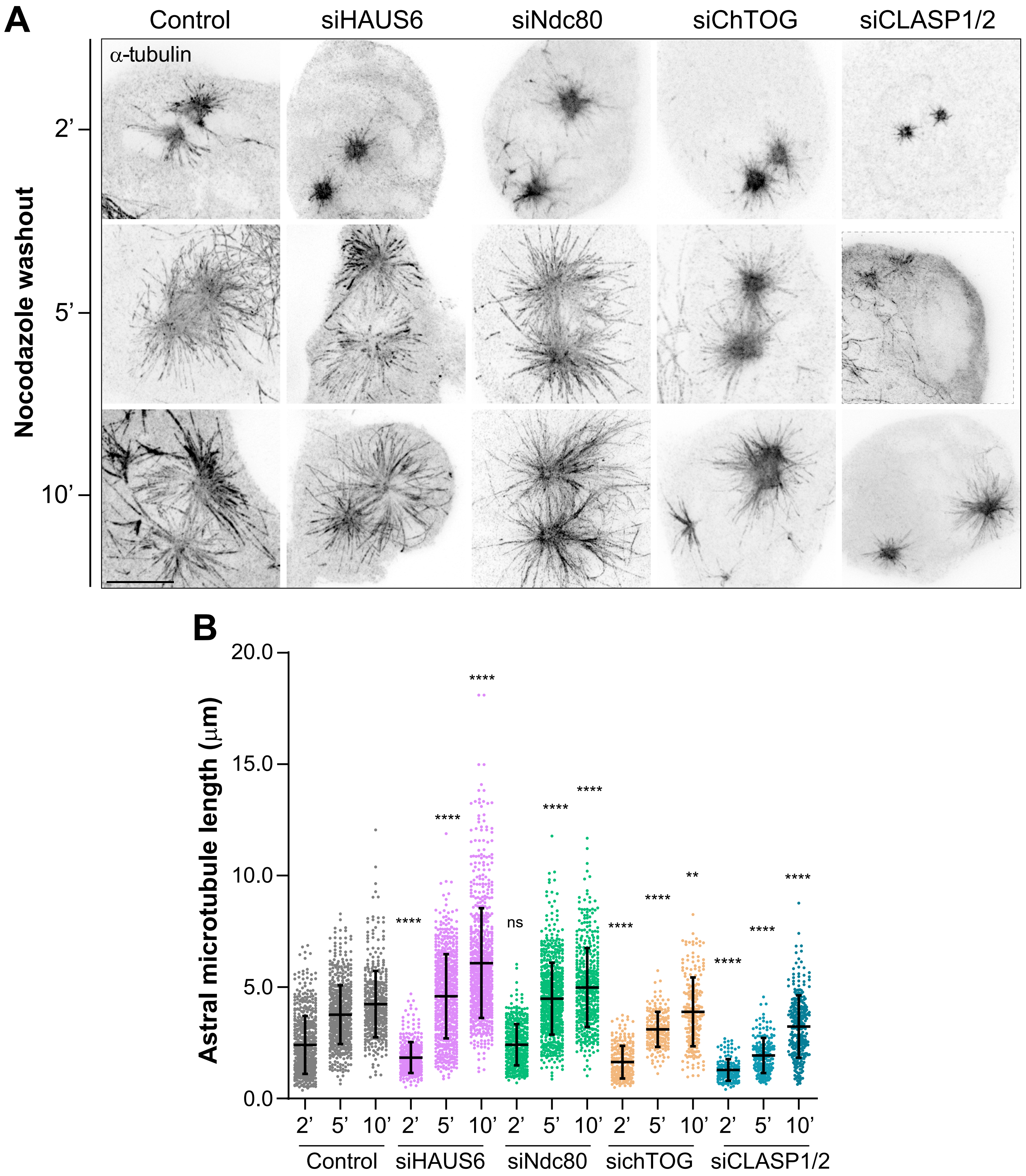


**Figure S4- Almeida et al.**

### Figure S5

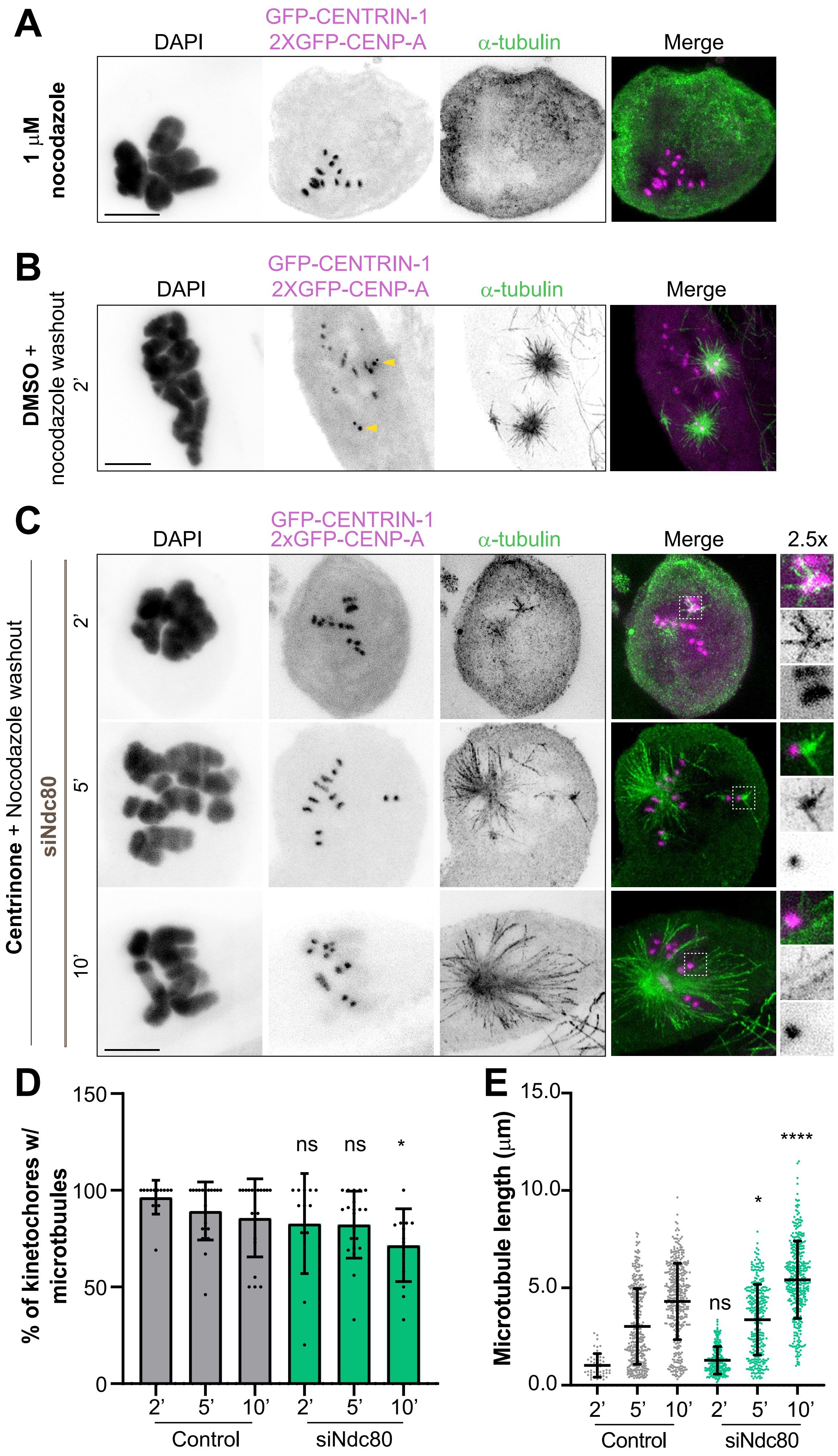


**Figure S5 - Almeida et al.**

### Figure S6

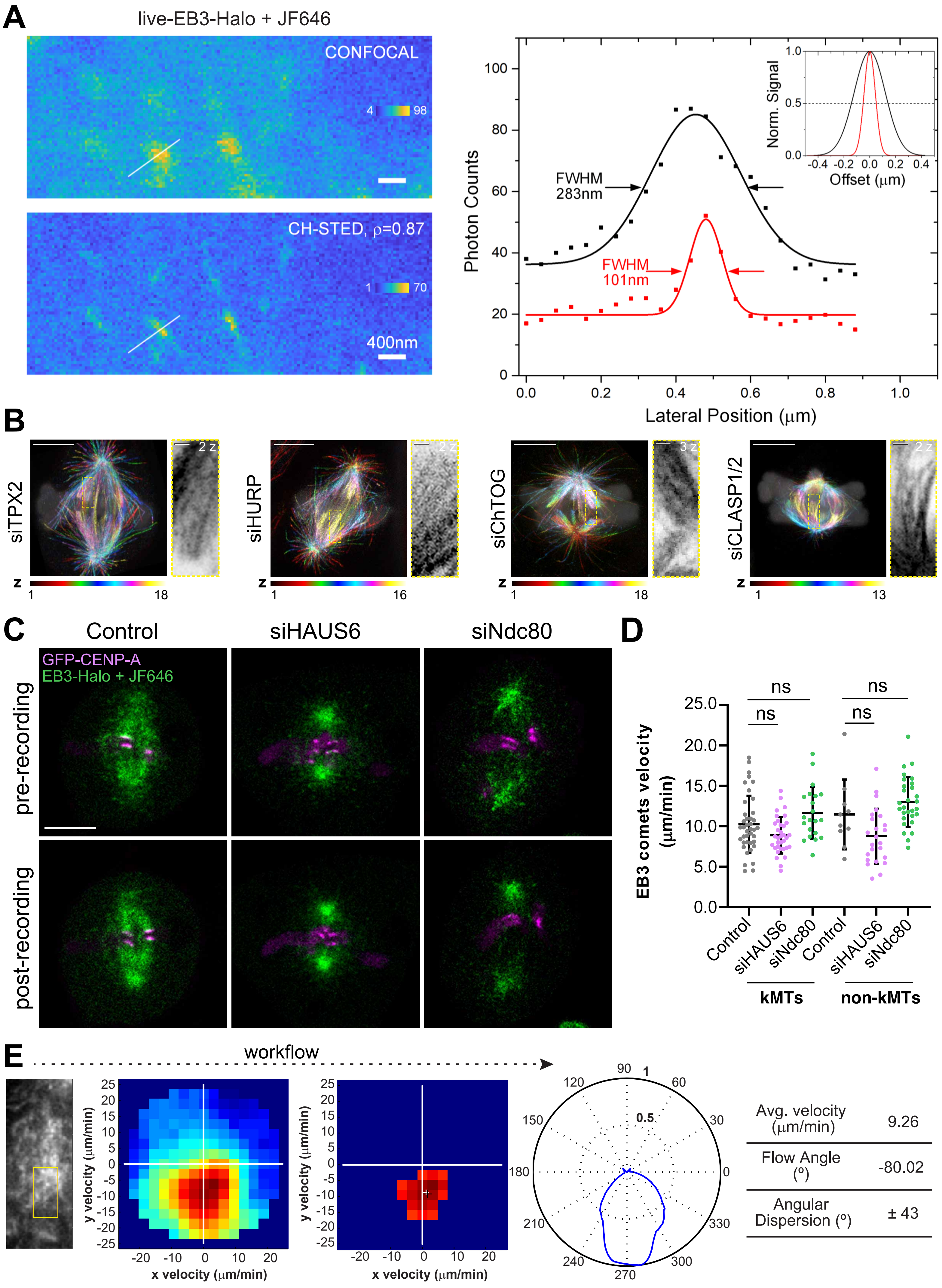


**Figure S6 - Almeida et al.**

### Figure S7

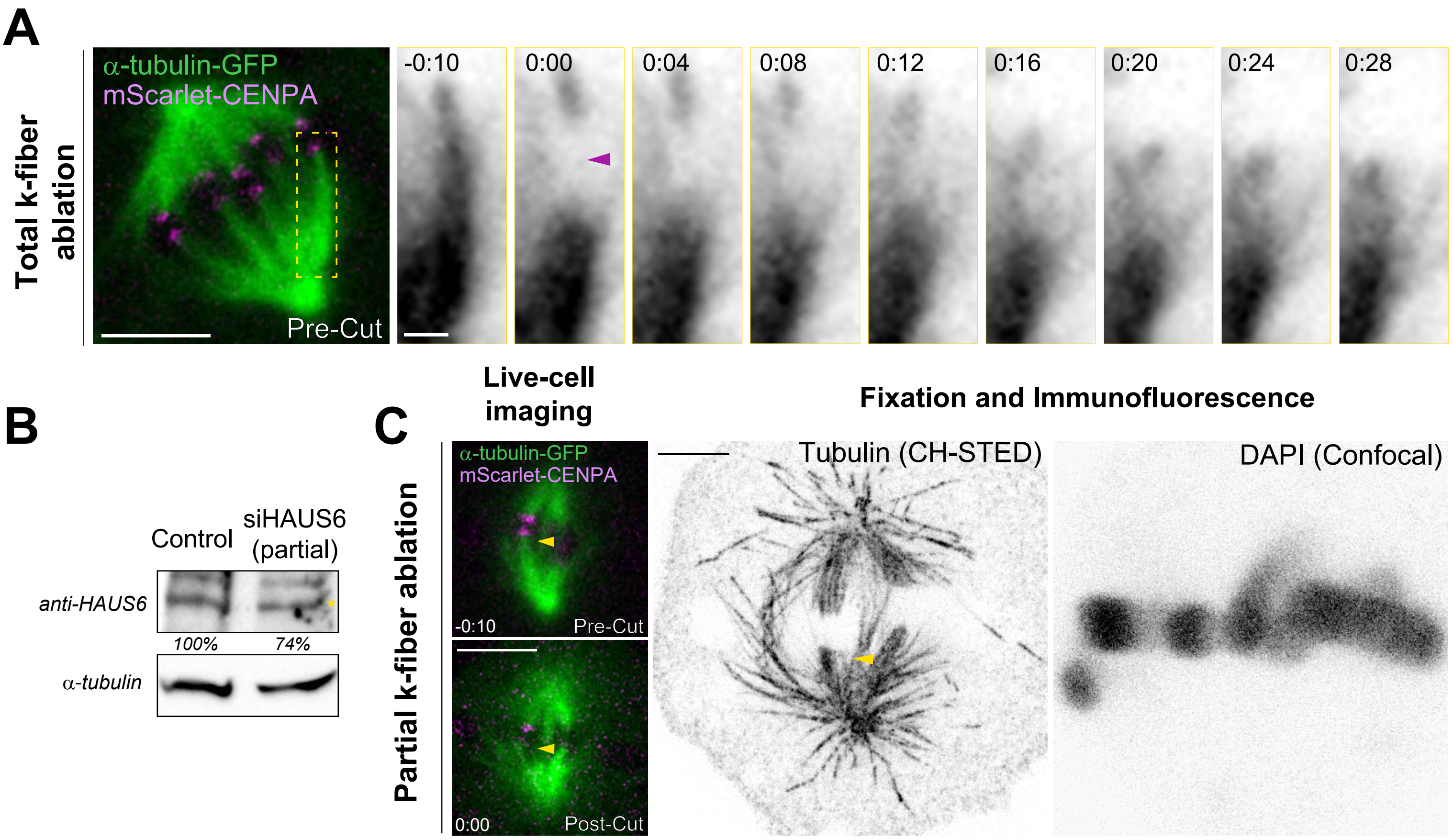


**Figure S7- Almeida et al.**
