## Supplementary material for "Kinetochore-mediated microtubule assembly and Augmin-dependent amplification drive k-fiber maturation in mammals": Table S1

**Table S1.** Antibodies used in this screening for immunoblotting and immunofluorescence.

| | Protein Name | Primary Antibody | | | |
|---|---|---|---|---|---|
| | | Source | Catalog Number/Reference | WB dilution | IF/STED dilution |
| 1 | MPS1 | Millipore (clone 3-472-1) | 05-682 | 1:1000 | |
| 2 | BUB3 | Gift from S. Taylor | (Lara-Gonzalez et al., 2012) | 1:1000 | |
| 3 | BUB1 | Gift from S. Taylor | (Taylor et al., 2001) | 1:500 | |
| 4 | BUBR1 | Abcam | Ab 200062 | 1:1000 | |
| 5 | MAD1 | Millipore | MABE867 | 1:1000 | |
| 6 | MAD2 | Bethyl | A300-300A | 1:500 | |
| 7 | CDC20 | Bethyl | A301-180A | 1:1000 | |
| 8 | TPR | Novus Biologicals | NB100-2867 | 1:1000 | |
| 9 | ChTOG | Santa Cruz Biotechnology | Anti-Ckap5 (H-4); sc-374394 | 1:500 | |
| 10 | CLASP1 | - | (Pereira et al., 2009) | 1:50 | |
| 11 | CLASP2 | - | (Pereira et al., 2009) | 1:10 | |
| 12 | CLIP-170 | Gift from N. Galjart Ab#2360 | (Molines et al., 2020) | 1:1000 | |
| 13 | EB1 | Abcam | 11B11 | 1:100 | |
| 14 | EB3 | Gift from A. Akhmanova | (Stepanova et al., 2003) | 1:3000 | |
| 15 | TTL | Proteintech | 66076-1-lg | 1:1000 | |
| 16 | HURP | Gift from P. Meraldi (R140) | (Silljé et al., 2006) | 1:200 | 1:500/1:100 |
| 17 | STMN1 | Proteintech | 11157-1-AP | 1:1000 | |
| 18 | ASTRIN | Gift from D. Compton | (Mack and Compton, 2001) | 1:500 | |
| 19 | HAUS6 | Gift from R. Uehara | (Uehara et al., 2009) | 1:500 | 1:500/1:50 |
| 20 | KTNB1 | Proteintech | 14969 | 1:1000 | |
| 21 | TPX2 | Proteintech | 11741-1-AP | 1:1000 | |
| 22 | Ndc80 | Abcam | ab3613 | 1:500 | |
| 23 | NUF2 | Abcam | ab3613 | 1:500 | |
| 24 | SPC24 | Abcam | ab3613 | 1:500 | |
| 25 | MIS12 | Gift from C. Conde | (Feijão et al., 2013) | 1:1000 | |
| 26 | SKA1 | Gift from P. Meraldi | clone-680 | 1:1000 | |
| 27 | SPINDLY | Gift from A. Desai | (Gassmann et al., 2010) | 1:1000 | |
| 28 | KNTC1 | Gift from Reto Gassman | (Gama et al., 2017) | 1:3500 | |
| 29 | CENP-I | Gift from P. Meraldi | (McClelland et al., 2007) | 1:250 | |
| 30 | CENP-F | BD Biosciences | 610768 | 1:1000 | |
| 31 | CENP-C | Gift from W. Earnshaw | (Saitoh et al., 1992) | 1.500 | |
| 32 | Aurora A | Novus Biologicals | NB100-267 | 1:1000 | |
| 33 | NINEIN | Gift from E. Nigg | (Logarinho et al., 2012) | 1:2000 | |
| 34 | PLK4 | Santa Cruz Biotechnology | Anti-Sak: sc-100413 | 1:100 | |
| 35 | Aurora B | Rockland (anti-AIM1) | 611082 | 1:500 | |
| 36 | Survivin | Novus Biologicals | NB500-201 | 1:1000 | |
| 37 | INCENP | Santa Cruz Biotechnology | sc-376514 | 1:250 | |
| 38 | NDEL1 | Abnova | H00054820-M01 | 1:1000 | |
| 39 | Dynactin | Gift from R. Gassmann | BD Transduction Lab. 610473 | 1:2500 | |
| 40 | NuMA | Gift from D. Compton | (Manning et al., 2007) | 1.2000 | |
| 41 | DYNEIN | ThermoFisher Scientific | PA5-49373 | 1:500 | |
| 42 | CENP-E | Santa Cruz Biotechnology | sc-376685 | 1:250 | |
| 43 | KIF20A | Bethyl Laboratories | A300-879A-M | 1:5000 | |
| 44 | KIF2A | Gift from D. Compton | (Ganem and Compton, 2004) | 1.10000 | |
| 45 | KIF2C | Gift from D. Compton | (Mack and Compton, 2001) | 1.500 | |
| 46 | KIF15 | Cytoskeleton Inc. | Cat#AKIN13 | 1:1000 | |
| 47 | KIF18A | Gift from AR. Maia | Bethyl Laboratories, A301-079A | 1:500 | |
| 48 | KIF18B | Gift from J. Welburn | (McHugh et al., 2018) | 1:1000 | |
| 49 | KIFC1 | Gift from D. Compton | sc-100947 | 1:1000 | |
| 50 | KIF11 | Sigma-Aldrich | HPA010568 | 1:1000 | |
| 51 | KIF23 | Proteintech | 28587-1-AP | 1:500 | |
| 52 | KIF4A | Thermo Fisher Scientific | pa5-30492 | 1:1000 | |
| 53 | KIF22 | Gift from S. Geley | 8612 | 1:500 | |
| 54 | SGO1 | Santa Cruz Biotechnology | sc-393993 | 1:500 | |
| 55 | CNDG2 | Gift from R. Oliveira (Novus Biological) | Citomed #NBP1-88202 | 1:1000 | |
| 57 | SECURIN | Thermo Scientific | 700791 | 1:500 | |
| 58 | SEPARASE | Santa Cruz Biotechnology | sc-390314 | 1.500 | |
| 59 | CDK1 | Santa Cruz Biotechnology | anti-cdc2; sc-54 | 1:500 | |
| 60 | CYCLIN-B1 | Cell Signaling | 4135 | 1:250 | |
| 61 | PRC1 | Santa Cruz Biotechnology | C-1; sc-376983 | 1:100 | |
| 62 | PLK1 | Abcam | ab115763 | 1:1000 | |
| 63 | VASH1 | Abcam | ab199732 | 1:250 | |
| 64 | VASH2 | ---- | --- | --- |  |

| | | | | | |
|---|---|---|---|---|---|
| 65 | PICH | Millipore | 04-1540 | 1:250 | |
| 66 | AlfatTubulin | Sigma Aldrich | B-5-1-2 | 1:10000 | 1:2000/1:200 |
| 67 | Vinculin | ThermoFisher Scientific | 700062 | 1:5000 | |
| 68 | GAPDH | Proteintech | 60004-1-Ig | 1:20000 | |
| 69 | CREST | Fitzgerald | #90C-CS1058 | | 1:2000/1:100 |
| 70 | Alfa-tubulin | Bio-Rad | MCA77G | ---- | 1:2000/1:100 |
| 71 | cMad2 | Santa Cruz Biotechnology | sc-65492 | | 1:250 |
| 72 | γ-tubulin | Sigma-Aldrich | Clone GTU-88 Mab #T6557 | | 1:5000/1:1000 |
